# Ablation of the choroid plexus attenuates hydrocephalus induced parenchymal edema but moderately reduces postnatal hippocampal neurogenesis

**DOI:** 10.64898/2026.03.30.714236

**Authors:** Aleksandr Taranov, Florian Mayrhofer, Sage Hamm, Eric Kniffen, Joshua Peter, Felissa Wallace, Olivia Lullmann, Lucas McClain, Olga V. Chechneva, Cameron Sadegh, Yu Luo

## Abstract

**Background:** Choroid plexus (ChP) produces cerebrospinal fluid (CSF) and regulates brain development and adult subventricular zone (SVZ) neurogenesis, but its role in postnatal hippocampal subgranular zone (SGZ) neurogenesis is not well understood. Infantile hydrocephalus is sometimes treated with partial lateral ventricle ChP cauterization, resolving clinical symptoms but largely preserving ventricular volumes. To understand the roles of the ChP/CSF system in normal conditions and the impact of ChP ablation or cauterization in hydrocephalus, novel tools for the manipulation of ChP-mediated CSF production are urgently needed.

**Methods:** We first tested the ROSA26-iDTR mouse model for ChP ablation at neonatal ages, then generated a novel AAV5-CMV-DTR vector with high ChP tropism to effectively reduce ventricular volume in neonatal mice. Genetic and gene therapy approaches were compared for their extent of ChP ablation, reduction of ventricular CSF volume assessed by MRI, and impact on postnatal hippocampal neurogenesis in neonates and adults. Lastly, we tested AAV-mediated ChP ablation in a kaolin model of hydrocephalus.

**Results:** In ROSA26-iDTR mice, ChP ablation and CSF volume reduction were robust at postnatal day (P)10-12, but ineffective at P3-5 with standard diphtheria toxin (Dtx) dosing (20 ng/g/day, 3 consecutive days). A higher dose (40 ng/g/day) at P3-5 mildly reduced CSF but caused neonatal mortality. In contrast, AAV5-CMV-DTR showed high tropism for ChP epithelium, producing near-complete loss of ventricular CSF in neonates. ChP/CSF loss in neonates or adult mice substantially reduced neuroblasts within the SVZ, but only moderately so in the SGZ, without altering the number of proliferating or apoptotic cells. In the kaolin model, AAV-mediated ChP ablation within 10 hrs after the onset of hydrocephalus profoundly reduced parenchymal edema and corpus callosum hyperintensity without causing ventricular collapse.

**Conclusions:** We demonstrate a role of the ChP/CSF in maintaining the neuroblast pool in both postnatal neurogenic niches. Although ROSA26-iDTR-mediated ChP ablation is inefficient before P10, the AAV5-DTR vector efficiently targets ChP and CSF production at both neonatal and adult ages, demonstrating the therapeutic potential of AAV-mediated ChP ablation for the treatment of hydrocephalus. The limited radiographic changes in the kaolin mouse model mirror the clinical findings following choroid plexus cauterization in humans.

## Introduction

The cerebrospinal fluid (CSF) produced by the choroid plexus (ChP) has long been recognized as a key regulator of brain development, homeostasis, and disease^1–3^. In addition to serving as a hydromechanical barrier and regulating electrolyte balance and waste clearance^4^, the CSF carries a rich repertoire of signaling factors mostly secreted by the ChP, including hormone carriers, cytokines, neuropeptides, and other bioactive molecules that may influence neurogenesis in neural stem cell (NSC) niches throughout life^5,6^. In the postnatal brain, the subventricular zone (SVZ) and the subgranular zone (SGZ) of the dentate gyrus represent two major neurogenic niches^7–10^ whose architecture and function are intimately shaped by their microenvironment, including signals derived from CSF^11–17^. In the process of neurogenesis, quiescent NSCs become activated and proliferate, giving rise to transiently amplifying progenitor cells, which then become neuroblasts (NBs). Newly born NBs migrate into the olfactory bulb from the SVZ, and deeper into the granular layer in the SGZ, where they ultimately differentiate into adult-born granule neurons^8,18^.

In contrast to embryonic neurogenesis, the cell-extrinsic molecular pathways regulating adult neurogenesis are relatively under-studied. In particular, understanding the role of the ChP/CSF mechanisms for regulation of endogenous self-renewal is critical, given the role of adult hippocampal neurogenesis in learning and memory^19–21^, stress-related disorders such as anxiety and depression^22–24^, epilepsy^25^, neurodegenerative disease^26^ and recovery from ischemic stroke^27–30^. New tools are urgently needed to better understand the influence of CSF composition and its circulation on these conditions.

We recently first discovered that the ROSA26-iDTR mouse line has an unexpected, Cre-independent DTR expression in the ChP epithelial cells, which makes them susceptible to diphtheria toxin (Dtx)-mediated cell death^31,32^. This enables noninvasive and temporally controlled ablation of the ChP and chronic loss of CSF volume in the adult brain^31^. Using this model, we demonstrated that ChP ablation in young adult mice (at the age of one month) causes rapid and stable depletion of CSF in all ventricles without overt motor or cognitive deficits at one month post-ablation. However, ChP/CSF loss resulted in a pronounced depletion of SVZ doublecortin-positive (DCX^+^) neuroblasts due to enhanced migration into the olfactory bulb, and markedly impaired ischemia-induced SVZ neurogenesis and neuroblast migration into the lesion site^31^. These findings established ChP/CSF as critical regulators of adult SVZ neuroblast migration and localization, and neuroreparative capacity following stroke.

Whether similar ChP/CSF-dependent mechanisms operate in the hippocampal neurogenic niche remains unknown. Adult hippocampal neurogenesis in the SGZ contributes to learning, memory, and affective behaviors^19,23^, epilepsy progression^33–35^, and declines with age^36,37^, yet the extent to which ChP-derived factors and CSF production support SGZ neurogenesis during the early postnatal and adult stages has not been assessed. Our prior work using ROSA26-iDTR achieved efficient ChP ablation and CSF loss in young adult and aged mice, but this tool has not been tested in early postnatal stages when SVZ and SGZ neurogenesis is highly active, nor in clinically relevant models of hydrocephalus, for which partial ChP ablation/cauterization is already a part of routine clinical practice in children under the age of 1 year^38^.

Hydrocephalus is a devastating neurological disorder characterized by excessive CSF accumulation, predominantly through impaired drainage due to congenital abnormalities or injuries (e.g., CNS infection, head trauma, neoplasms or intracranial hemorrhage) and less commonly by CSF overproduction (e.g., ChP carcinomas of the lateral ventricles)^39,40^. Hydrocephalus typically results in ventricle expansion and compression of surrounding brain parenchyma, and is often associated with peri- ventricular T2 signal change on MR imaging^41–43^. Transependymal flow is indistinguishable from parenchymal edema, which is associated with white matter demyelination, axonal injury and neuroinflammatory gliosis ^44,45^. Despite the high burden of disease, current interventions for hydrocephalus, such as surgical ChP cauterization^38^, endoscopic third ventriculostomy^38,46,47^, and CSF shunting are characterized by high rates of complications. Therefore, novel minimally invasive one-time treatments for hydrocephalus are urgently needed. CSF-mediated ChP-targeted gene therapy avoids the toxicity of systemic intravenous vector delivery and represents a promising, nascent strategy to both study the ChP/CSF system and to treat hydrocephalus and other CNS disorders^48–50^.

In the present study, we first ask whether ChP/CSF loss in young adult mice affects postnatal SGZ neurogenesis, as it does for the SVZ. Second, we extend the toolkit for ChP ablation by developing an adeno-associated viral (AAV2/5) DTR-based tool for selective ChP ablation in neonatal and adult mice, enabling a broader age range for ChP ablation, as compared to ROSA26-iDTR-mediated ChP ablation. Third, we demonstrate the use of AAV-mediated ChP ablation as a proof-of-principle approach for the treatment of hydrocephalus.

We show that ChP/CSF loss in young adult ROSA26-iDTR mice leads to a moderate reduction of SGZ DCX^+^ neuroblasts without altering proliferation and apoptosis in the SGZ, suggesting a role for ChP/CSF in supporting postnatal hippocampal granule cell differentiation and/or survival. We further demonstrate that ROSA26-iDTR-mediated ChP ablation is inefficient before postnatal day 10 (P10), and that, in contrast, an AAV5-CMV-DTR viral approach provides an effective and highly specific method to ablate ChP epithelial cells and reduce CSF volume when initiated at P1, allowing for the study of ChP/CSF functions in early postnatal development. Using this viral approach in wild-type mice, we find that neonatal ChP/CSF loss decreases SVZ DCX+ NBs and leads to clustered neuroblasts in the SVZ lateral wall, accompanied by reorganization of ependymal cells around these clusters. Additionally, neonatal ChP/CSF loss moderately reduces SGZ DCX^+^ cells and hippocampal volume, again without a difference in Ki67+ proliferating cells or TUNEL^+^ apoptotic cells at P30. In the kaolin model of experimental hydrocephalus, we show that ChP ablation at an acute stage (6-10 h) after hydrocephalus induction reduces parenchymal edema and corpus callosum hyperintensity seen on day 4, which substantially expands at day 7 post-induction in control AAV-treated hydrocephalus mice but not in ChP-ablated mice.

Together, these findings extend the cell-extrinsic regulatory role of the ChP/CSF axis to both sites of adult neurogenesis and establish complementary genetic and viral tools to manipulate ChP/CSF across postnatal development and adulthood. Moreover, we demonstrate the potential therapeutic utility of this tool as treatment for hydrocephalus.

## Materials and Methods

### Animals

All animal experiments, husbandry procedures, and surgical interventions were conducted in accordance with the protocols approved by the University of Cincinnati and University of California Davis Institutional Animal Care and Use Committees (IACUC) and fully complied with the guidelines established by the National Institutes of Health (NIH) Guide for the Care and Use of Laboratory Animals. All mice were maintained on a C57BL/6J background. Wildtype C57BL/6J (Jax #000664) and ROSA26iDTR mice (Jax # 007900) were obtained from Jackson Laboratories and bred in-house. The mice were housed in the animal facility of University of Cincinnati or University of California Davis on a 12-h light/12-h dark diurnal cycle. Food and water were provided *ad libitum*. The age of mice in each experiment is indicated in individual figure legends. Both female and male mice were used in this study, and the sex of each mouse is labeled in the figures using circles (female) or triangles (male) for each data point.

### Administration of Diphtheria toxin (Dtx) to ablate the choroid plexus (ChP) in the iDTR mice

Recombinant *Corynebacterium diphtheriae* toxin was obtained through Sigma (cat. # D0564-1MG or 322326, similar results) as a lyophilized powder and reconstituted in 0.9% saline at 2 mg/ml. The stock solution was stored at - 80 °C and diluted with ice-cold saline as needed. The resulting working solution was kept on ice and used within one hour after preparation. As described previously^31^, Dtx was administered for three consecutive days with no more than 24 h between injections. Adult mice received Dtx i.p. from 2 ng/ul working solution at 20 ng/g/day. Neonatal mice received Dtx subcutaneously at 5-10 ng/ul, depending on the dose, for three days (regimens and doses described in figures and figure legends).

### Viral Vector Design and Production

To achieve highly specific transduction of the ChP epithelium in early postnatal mice, recombinant Adeno-Associated Virus serotype 5 (AAV5) vectors were utilized. Two distinct viral constructs were acquired from VectorBuilder: control AAV5-CMV-eGFP:WPRE, and AAV5-CMV-DTR:WPRE.

### Tissue Collection for Immunohistology

All animals were euthanized by administration of avertin (2.5%) followed by transcardial perfusion with ice-cold 0.1 M phosphate buffer (pH 7.2) and then with 4% paraformaldehyde in 0.1 M phosphate buffer (pH 7.2). The brain was dissected out and post-fixed in 4% paraformaldehyde overnight at 4 °C and sequentially dehydrated in 20% and 30% sucrose in 0.1 M phosphate buffer (pH 7.2) for cryoprotection.

### Immunostaining

#### Free-floating IF staining of coronal sections or whole ChP tissues

For free-floating immunofluorescent staining, the brain tissue was frozen in OCT on dry ice and sectioned at 30 μm on a cryostat. Whole dissected ChP tissue or coronal brain sections were washed 3x and blocked in blocking buffer (4% BSA, 0.3% Triton X- 100 (Acros Organics) in 0.1 M phosphate buffer, pH 7.2) for 1 hr at RT with shaking and transferred into primary antibody solution in blocking buffer for incubation overnight at 4 °C on a shaker. Then, sections were washed in 1XPB before incubation in secondary antibodies (conjugated with Alexa 488, Alexa 555, Alexa 647, Alexa 790 1:500; Life Technologies, Carlsbad, CA, USA or Jackson Immuno Research, West Grove, PA, USA) dissolved in blocking buffer for 4 hours at RT with shaking. For the ChP whole mounts, the procedure was the same except for the secondary antibody incubation overnight at 4 °C with shaking. Free-floating sections were mounted onto microscope slides (12-550- 15, Fisherbrand) and coverslipped in Mowiol 4-88 (17951, Polysciences Inc.) mounting medium.

#### Staining of Directly Mounted Coronal Cryosections

For ChP area quantification using coronal brain sections, brain tissue was sectioned coronally at 40 µm into 4 sets on a cryostat and directly mounted onto glass slides to prevent potential loss of ChP tissue during free-floating staining. Slides were washed 3x and blocked for 1 hour at RT in blocking buffer using CoverWell incubation chambers (645502, Grace Bio-labs), 950 µl per chamber. Then, the slides were incubated for 72 hours (to ensure good penetration) at 4 °C in primary antibody diluted in blocking buffer, washed for 30 min (repeated 3 times) and then incubated in secondary antibody diluted in blocking buffer for 48 hours at 4°C. All incubations were done in CoverWell chambers.

#### SVZ whole-mount staining for characterization of neurogenesis and ependymal cells

Brain tissue was harvested after 4% PFA perfusion. SVZ whole-mounts were dissected as described^31^ and washed 3×10 min in 0.1% Triton X-100 in 0.1 M PB. Then, the mounts were blocked in 4% BSA in 0.1 M PB (pH 7.2) with 2.3% Triton X-100 for 1 hour at RT. The whole mounts were transferred into the primary antibody solution and incubated for 48 hours at 4 °C with shaking. Then, the tissue was washed 3×30 min in 0.1% Triton X-100 in 0.1 M PB and transferred into secondary antibody solution and incubated for 48 hours at 4 °C with shaking, washed 3 × 30 min in 0.1% Triton X-100 in 0.1 M PB, mounted on microscope slides and coverslipped.

#### Image Acquisition

Confocal imaging of stained brain sections was carried out using a Leica Stellaris 8 confocal microscope. Quantification of cell counts was carried out using ImageJ software. At least 3 coronal sections and up to 6 sections with respective regions of interest were quantified for each mouse and averaged to a single data point in data analysis.

Machine learning-assisted segmentation and cell counting were performed using an online analysis tool from www.biodock.ai. The classifier was trained on a set of single-channel DCX representative images from multiple animals. Training images were uploaded, and a class was assigned so that DCX-positive cells could be detected. Labels were manually created and assigned to this class during the training phase. Results were evaluated, and the model was retrained for 3 iterations to improve accuracy. Following this, the entire dataset was segmented and counted by the model. Randomly selected images were also quantified manually, showing excellent agreement with automatic quantification.

#### TUNEL Staining in the SGZ

Apoptosis was evaluated in tissue cryosections using the ApopTag® Fluorescein in situ Apoptosis Detection Kit (Millipore, Cat# S7110). Fixed sections were washed in PB buffer and then permeabilized in pre-cooled ethanol: acetic acid (2:1, v:v) for 5 minutes at -20 °C. Tissue was then treated with Equilibration Buffer for 10 seconds at room temperature. Sections were then incubated with working-strength TdT enzyme in a humidified chamber at 37 °C for 1 hour. The reaction was then terminated in Stop/Wash buffer for 10 minutes. Following this, co-staining of Ki67, DCX, and DAPI was done according to the protocol for directly mounted sections. Following primary and secondary antibody staining, the sections were incubated with a working strength anti-digoxigenin-fluorescein conjugate for 30 minutes at room temperature in the dark.

#### Primary Antibodies

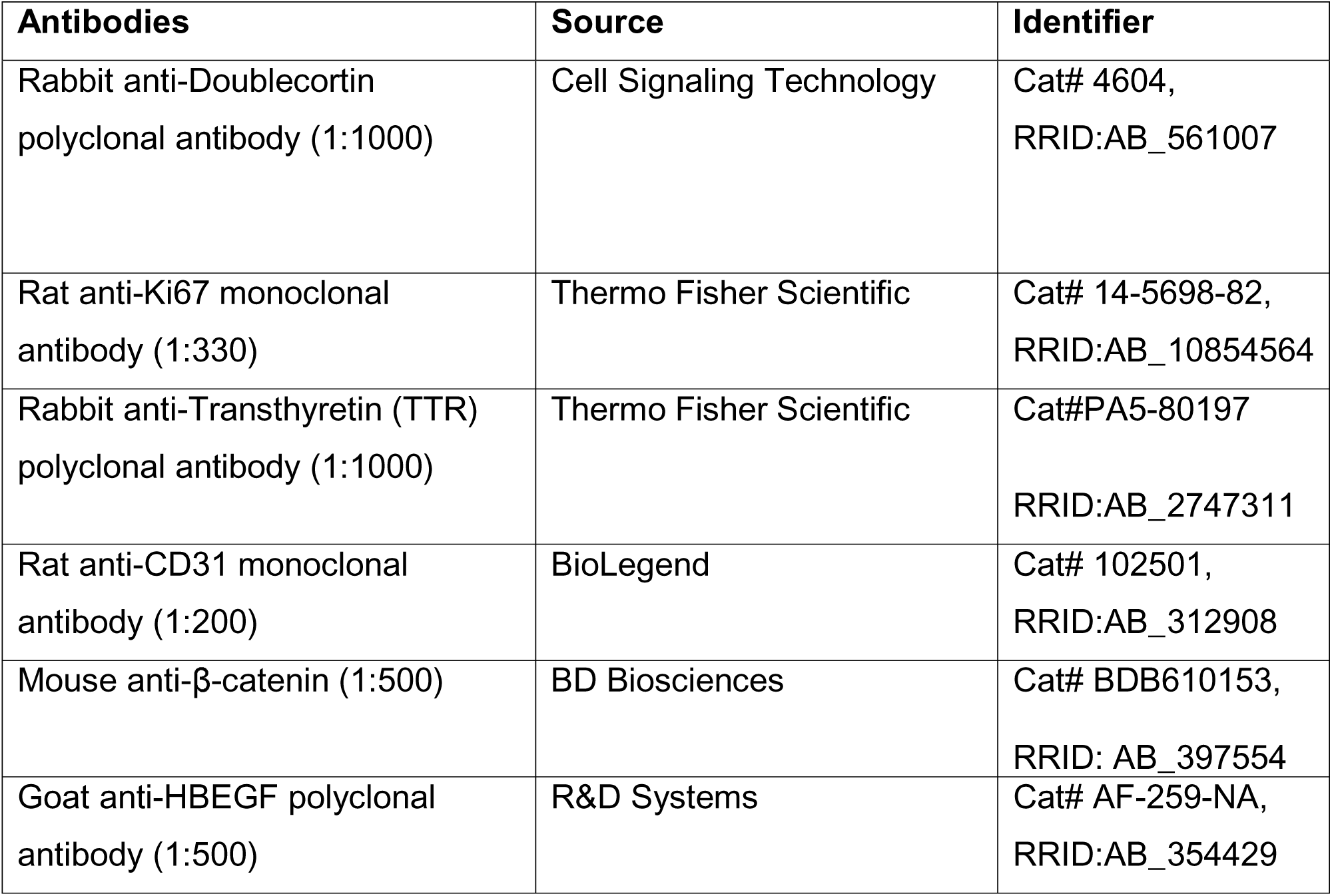

| Antibodies | Source | Identifier |
| --- | --- | --- |
| Rabbit anti-Doublecortin polyclonal antibody (1:1000) | Cell Signaling Technology | Cat# 4604,<br>RRID:AB_561007 |
| Rat anti-Ki67 monoclonal antibody (1:330) | Thermo Fisher Scientific | Cat# 14-5698-82,<br>RRID:AB_10854564 |
| Rabbit anti-Transthyretin (TTR) polyclonal antibody (1:1000) | Thermo Fisher Scientific | Cat#PA5-80197<br>RRID:AB_2747311 |
| Rat anti-CD31 monoclonal antibody (1:200) | BioLegend | Cat# 102501,<br>RRID:AB_312908 |
| Mouse anti-β-catenin (1:500) | BD Biosciences | Cat# BDB610153,<br>RRID: AB_397554 |
| Goat anti-HBEGF polyclonal antibody (1:500) | R&D Systems | Cat# AF-259-NA,<br>RRID:AB_354429 |

### Stereotaxic Viral Injections and Surgical Procedures

#### Neonatal Intracerebroventricular (ICV) Injections

As described previously^51^, P1 C57Bl/6J pups were cryoanesthetized and injected with 2 µl of AAV5 CMV-eGFP control (1.75X 10^12^ GC/mL) or CMV-DTR (1.41X 10^12^ GC/mL) into the left lateral ventricle. The needle was inserted through the skull at 40% of the length between the eye and lambda at 3 mm deep, and the virus solution was infused over the course of 1 minute, and the needle was left in place for 30 seconds to prevent backflow. The pups were allowed to recover in an incubator and returned to the nest once locomotion resumed.

#### Adult Stereotaxic ICV Injections

Adult C56Bl/6J mice were anesthetized with isoflurane (3% induction, 1.5% maintenance) and placed into a stereotaxic frame. Buprenex ER analgesia was administered subcutaneously. The skull was exposed, and a burr hole was drilled at AP-0.35 mm, ML: -1.0 mm. A 33g Hamilton syringe was lowered to 2.5 mm, and 6 µl of CMV-eGFP (2.1 × 10^12 GC/mL) or 4 µl of CMV-DTR (3.03 × 10^12 GC/mL) of virus solution was infused at 1 µl /minute. Then, the needle was slowly withdrawn 1 mm and left in place for 10 minutes to prevent backflow. After complete withdrawal, the incision was sutured, and the animal was allowed to recover in a heated incubator.

#### Kaolin-induced hydrocephalus

Male C57BL/6J mice (8–12 weeks old) were used in this experiment. Each mouse was deeply anesthetized with isoflurane (3–5%) until unresponsive to noxious stimulation and secured in a stereotaxic frame (Kopf Instruments). Throughout the surgical procedure, animals were maintained on a warming blanket covered with a Chux pad. Ophthalmic lubricant was applied to both eyes to prevent corneal drying. The NSAID carprofen (5 mg/kg) was administered subcutaneously at the scruff of the neck immediately after anesthesia induction and prior to the skin incision. Hair was removed from the back of the head using Nair, and the surgical site was prepared with alternating applications of betadine and 70% alcohol. A 1-cm midline vertical skin incision was made over the back of the neck, extending from the occiput to the first cervical vertebra (C1). The underlying muscle layers were bluntly separated and divided along the natural midline plane to minimize tissue damage. A 5 µL suspension of kaolin (250 mg/mL in PBS; Sigma) was injected into the cisterna magna at a rate of 1 µL/min using a 30- gauge needle attached to a 100 µL gas-tight Hamilton glass syringe. Following the injection, the paraspinal muscles were allowed to return to their anatomical position, and the skin incision was closed using Vetbond tissue adhesive. The mice were allowed to recover in a warmed cage. Postoperative analgesia was provided with subcutaneous carprofen (5 mg/kg) administered at the scruff of the neck (distal to the incision) 24 and 48 hours after surgery.

### T2-weighted MRI, brain ventricular measurement using T2-weighted MRI anatomical scans, and 3D MRI reconstruction (University of Cincinnati)

MRI studies (except for kaolin experiment, which is described separately below) were performed on a vertical wide-bore 9.4T Bruker Avance III HD scanner with a 36mm proton volume coil. Mice were anesthetized with 1.5 - 2% isoflurane and kept warm with circulating air. Temperature and respiration rate were monitored with equipment from Small Animal Instruments, Inc. (SAI, Inc., NY). T2-weighted anatomical coronal images of the brain were acquired with a fat suppressed two-dimensional (2D) rapid acquisition with relaxation enhancement (RARE) sequence using the following parameters: TR 4 sec, TE 71.5 ms, echo spacing 6.5 ms, 9 or 15 slices, slice thickness/gap 0.75/0.3 mm, RARE factor 20, receiver bandwidth 67k, averages 4, matrix 192×192, FOV 28.4×28.4 mm, and total scan time 2:24 minutes. Ventricular volumes were quantified by outlining and measuring ventricular space for each animal from the individual T2-weighted coronal images (sections ranging from Bregma AP: +1.5 mm to AP: − 4 mm) using ImageJ. CSF/ventricular volume was calculated by multiplying the area of ventricular space by the thickness of each slice (mm) of the acquisition. For 3D MRI reconstructions, fluid-sensitive images of the brain were acquired with a fat suppressed three-dimensional (3D) RARE sequence using the following parameters: TR 2 sec, TE 275 ms, echo spacing 11.5 ms, RARE factor 60, receiver bandwidth 104k, averages 4, matrix 320×108×80, FOV 48×16.2×12mm, and total scan time 7 minutes 44 seconds. To reconstruct ventricular spaces in 3D and quantify their volumes, images were imported into Imaris 10 and 3D ventricular surfaces were reconstructed from a series of DICOM files. The surfaces were generated using the Surfaces tool with a surface detail of 0.1 and seed point diameter of 0.8. The resulting 3D ventricular surfaces were manually divided into separate ventricles and parenchymal edema surfaces where appropriate. The volumes were quantified using the Measurements tool in Imaris.

### T2-weighted MRI fluid sensitive and anatomical scans for Kaolin experiment (UC, Davis)

For Kaolin experiment at UC Davis, MRI was performed using a 7-T Bruker BioSpec scanner equipped with a RF ARR 300 1H M coil (Bruker) coil. Mice were anesthetized with 1.5–2.0% isoflurane in oxygen and maintained at 37°C throughout imaging. T2- weighted images were acquired using a rapid acquisition with relaxation enhancement (RARE) sequence with the following parameters: repetition time (TR) = 11,200 ms, echo time (TE) = 160 ms, RARE factor = 16, field of view = 24 × 24 mm^2^, acquisition matrix = 200 x 192, 40 slices with 0.3 mm slice thickness, yielding an in-plane spatial resolution of 0.12 × 0.125 mm and a voxel size of 0.12 × 0.125 × 0.30 mm³. The total acquisition time was 8 minutes and 57 seconds. For anatomical scans, T2-weighted images were acquired using a RARE sequence (TR = 4,000 ms, TE = 67.2 ms, RARE factor = 12) with a field of view of 24 × 24 mm², an acquisition matrix of 200 × 192, yielding an in-plane spatial resolution of 0.120 × 0.125 mm². Twelve contiguous 1 mm thick slices were acquired, resulting in a voxel size of 0.120 × 0.125 × 1.0 mm³. The total acquisition time was 4 minutes and 16 seconds. Images were reconstructed using ParaVision4 software and analyzed in Imaris software. For post-kaolin corpus callosum hyperintensity quantification, four consistently located T2-weighted coronal slices were analyzed in ImageJ. The corpus callosum was manually traced, and the intensity and integrated density were measured from the resulting ROI.

### Open Field Test

The open field apparatus (Omnitech electronics Inc, Columbus, OH) consisted of 2 sets of 8 photobeam arrays for animal horizontal activity detection and 1 set of 8 photobeam arrays for vertical activity detection. Each monitor was installed on a cube frame inside of a closed cabinet. A 42Lx42Wx31T cm Plexiglas box divided into 4 chambers to allow for simultaneous recording of two animals was placed inside of each frame. Mouse bedding was evenly placed on the bottom of the box. Neonatal mice that received AAV5-eGFP or AAV5-DTR at P1 were placed in the chambers for 1h to monitor novelty-induced locomotor activity at P30. The chambers were washed using a steam mouse cage washer between tests. Animal activity was tracked using automated Fusion software (Omnitech electronics).

### Quantification and Statistical Analysis

All data were analyzed using SigmaPlot 12.0. Results are expressed as mean ± SEM. Each data point represents a single animal. Only one outlier was identified in all the data sets using the ROUT method with Q=1% in Prism. The outlier data point is from Fig 5F, one of the AAV-DTR+Dtx mouse (orange points), is ablated less effective than the rest of the group, which we did not remove. Statistical analysis was performed using the two-sided Student’s t-test, two-sided Welch’s t-test, two-sided Mann-Whitney U test, and two-sided one- or two-way analysis of variance (ANOVA), or Kruskal-Wallis test, as appropriate, with Tukey’s or Dunn’s post-hoc tests. When the data did not meet the assumptions of normality or equal variance, the corresponding nonparametric tests were used. The significance level was set at α = 0.05. The number of animals used in each experiment was plotted as a single data point and is indicated in Figure legends. Graphs were made in GraphPad Prism 10.6.1. Exact p-values are indicated in all figures or indicated as n.s when p ≥ 0.05.

## Results

We have previously shown that ChP ablation at P30 (using iDTR mice treated with Dtx) results in a profound decrease in the pool of DCX+ CSF-contacting NBs that line the lateral ventricle wall at the SVZ^31^. In this study, we investigate the role of ChP/CSF in regulating hippocampal neurogenesis, which occurs in a niche in proximity to both the lateral and third ventricles, though not traditionally considered to be regulated by the ChP/CSF.

First, we confirmed a robust ChP ablation phenotype using the iDTR mice (treated at 1 month of age) with a dosage of 20ng/g/day for three consecutive days (abbreviated as a multiplicative of 3, or “X3”) and validated substantial loss of CSF volume in all ventricles (Fig 1 A-D), consistent with our previous report^31^. Mice were harvested at 3 months post-ablation, a time point when we previously observed a pronounced decrease in SVZ DCX+ neuroblasts. We quantified SGZ DCX+ neuroblasts and Ki67+ proliferating cells and found that ChP ablation induces a significant decrease in the numbers of DCX+ cells (Fig. 1E-G, 18% decrease) in the SGZ without affecting proliferating Ki67+ cells (Fig 1H-L). These results show that the loss of ChP/CSF in adult iDTR mice decreased postnatal neurogenesis but not cell proliferation in SGZ, suggesting that ChP/CSF may be required for the maintenance of the neuroblast pool at the SGZ, in a similar manner to the SVZ.

**Fig 1.**
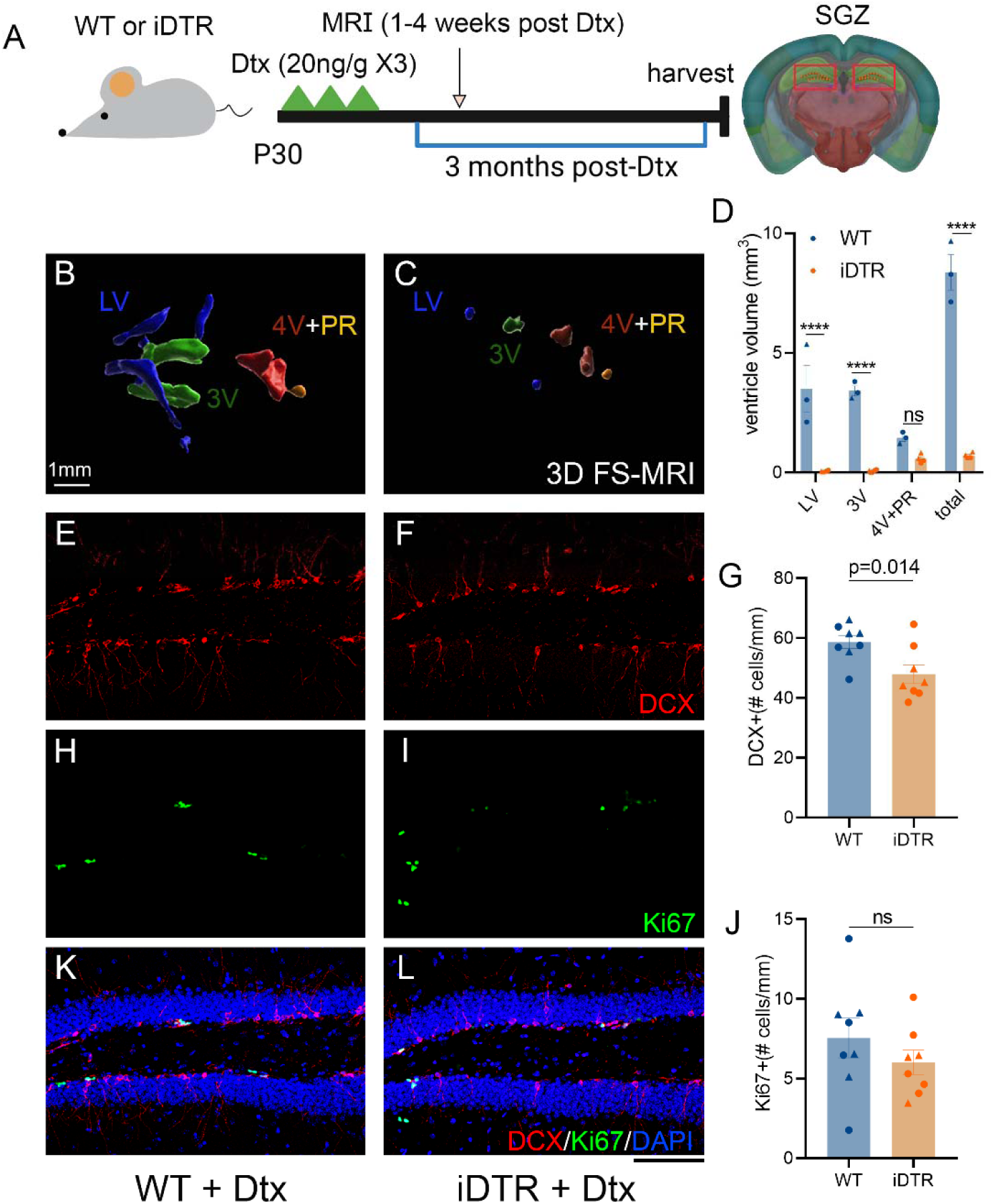
ChP ablation in adolescent ROSA26-iDTR mice leads to decreased ventricular volume and SGZ neurogenesis. (A) The experimental timeline. (B-C) Reductions in ventricular volume shown by representative 3D reconstruction of Fluid-sensitive MRI (B) in WT mice or (C) iDTR mice treated with 20ng/gX3 Dtx. (D) Quantification of ventricular volumes. (E-L) Representative images of DCX, Ki67 and DAPI staining in experimental groups. (G) Quantification of DCX+ cells in SGZ and (J) Quantification of proliferating cell numbers (Ki67+). Mean±SEM. Each data point is the data from an individual animal (average of 3-6 brain sections for IHC). Circle= female and Triangle=male. ****, *p*<0.0001 Two-Way ANOVA, for panel (D), or *p*=0.014 Student’s t-test for panel G or ns= not significant for panel J. Scale bar = 100 µm or as indicated.

Given this finding of ChP-dependent maintenance of the adult SGZ neuroblast pool, we hypothesized that the deleterious effect of ChP ablation might be more pronounced at earlier stages of development where reductions in hippocampal neurogenesis can contribute to epilepsy^33^ and affective disorders^52,53^. To investigate the role of the ChP in early postnatal neurogenesis, we next tested ChP ablation in P3-P5 mice using either 20 or 40ng doses of Dtx administered i.p. on 3 consecutive days (abbreviated as “20ng/gX3” or “40ng/gX3”; Fig. 2 A). Mice underwent 3D fluid-sensitive and T2 anatomical MRI on postnatal day 30 (Fig. 2 B, Fig. S1). Our data show that Dtx treatment in neonatal iDTR mice produces a much smaller CSF volume reduction compared to adult Dtx-treated mice. The 20ng/g daily treatment from P3-5 did not alter the total volume of all ventricles, while 40 ng/g daily treatment from P3-5 only showed modest CSF volume reduction, which is significant for the 3V and total CSF volume (LV+ 3V and PR+4V as a total measurement) but was not significant for other ventricle individually (Fig. 2 C). The high dosage of Dtx is likely to produce undesirable off-target effects, as reported previously^54–56^. Indeed, we find considerable mortality by P15 in both iDTR or WT neonates treated with 40 ng/g/day of Dtx, but not 20 ng/g/day of Dtx (Fig. S2 A, B), which limits the utility of this genetic model for ablation of the neonatal ChP.

**Fig 2.**
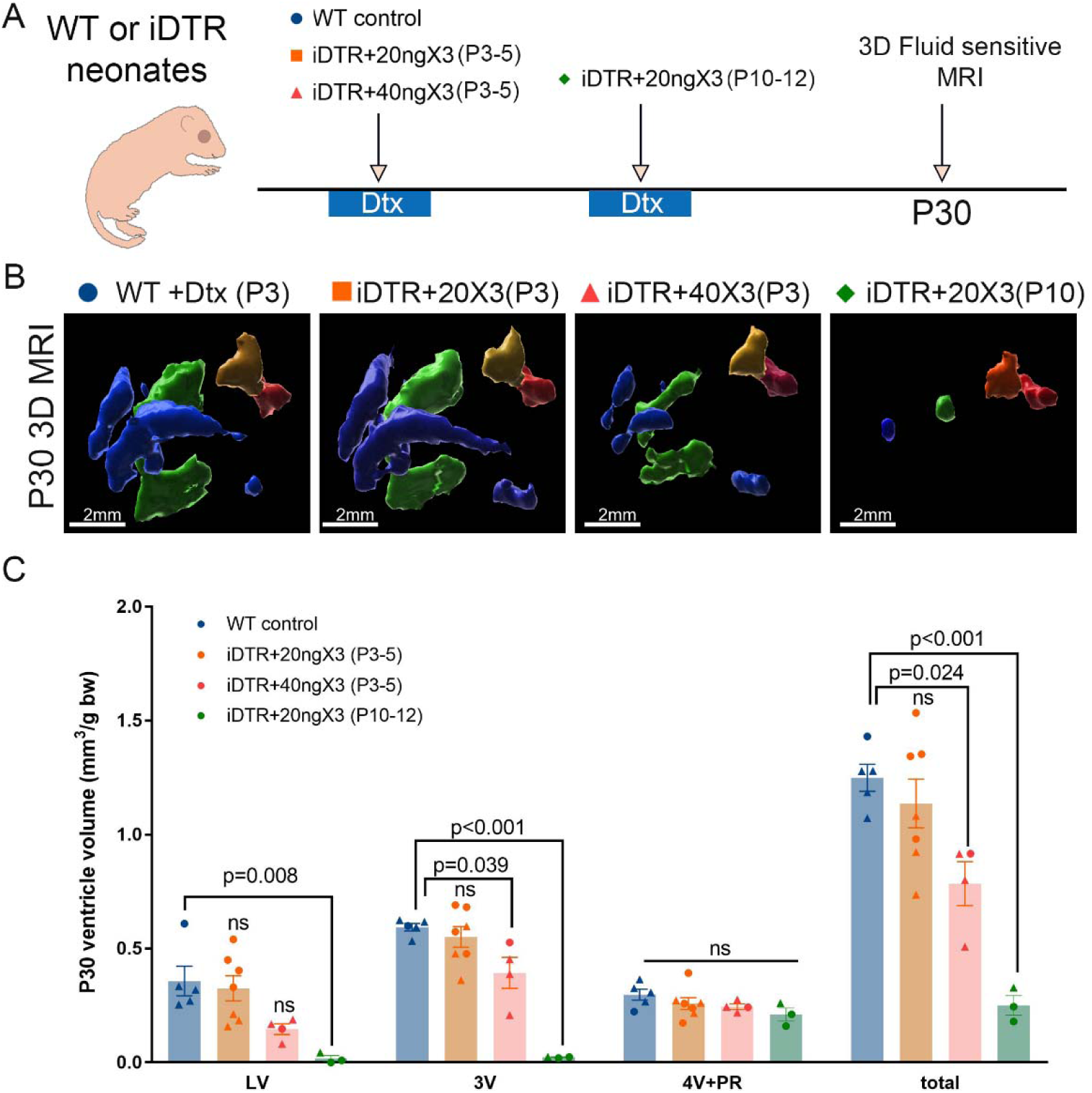
ROSA26-iDTR model does not efficiently ablate ChP or reduce CSF volume in neonates before P10. (A) Experimental timeline and description of the four groups of mice (genotype and Dtx treatment dosage/timing of treatment). (B) Representative 3D Imaris-reconstructed ventricular volume in the four groups of mice. (C) Quantification of individual ventricles and total ventricular volume. Mean±SEM. Each data point is the data from an individual animal. Circle= female and Triangle=male. *p-* value as indicated, one-way ANOVA for each ventricle for panel C.

The limited effect of Dtx at neonatal ages may be due to relatively low expression levels of the “leaky” DTR gene in postnatal ChP development. We hypothesize that the levels of DTR in ChP subsequently increases with developmental age, making Dtx treatment more efficient at later timepoints. Indeed, at P10-12, the low dose Dtx (20ng/g daily) was able to substantially reduce CSF volume in all ventricles (Fig 2C and supplemental Fig 1), similar to the level of CSF volume loss we observed in adult iDTR mice. This result shows that although the iDTR mouse line is an effective tool for ChP ablation in mice that are older than P10, an alternative strategy for ChP ablation is necessary for earlier timepoints in development. Such a tool would allow the study of early hippocampal neurogenesis and enable the potential therapeutic application to CSF disorders such as hydrocephalus.

To develop a more flexible and therapeutically relevant method of ChP ablation at any age, we designed the AAV5-CMV-DTR viral vector expressing the simian heparin-binding EGF-like growth factor (HB-EGF or DTR^57^) under the cytomegalovirus (CMV) promoter. Previous studies have shown that the AAV5 serotype can be effectively used for embryonic or postnatal ChP targeting^49,50,58^ and shows selective and strong transduction of the ChP epithelium without the need for ChP-specific promoters.

We first tested the efficiency and specificity of the AAV5 serotype in neonatal mice using the AAV5-CMV-eGFP viral vector. AAV5 CMV-eGFP viral vector was injected unilaterally into the lateral ventricle of neonatal P1 pups as previously described^51^ (Fig. 3A), and ChP whole mounts or coronal sections were analyzed for eGFP and TTR colocalization. Our data demonstrate that the AAV5-CMV-eGFP transduces ChP epithelial cells (CPECs) with high efficiency and specificity (Fig. 3B). We only observed sparse eGFP-positive cells outside of ChP (Fig 3B). In contrast, we show a high percentage of eGFP+ ChP epithelial cells (completely colocalizing with TTR, a marker of CPECs) in LV ChP (Fig 3C), 3V ChP (Fig 3D), and 4V ChP (Fig 3E), (> 70% TTR-positive cells eGFP+, Fig 3F). Immunostaining of ChP whole mounts shows similar results for LV ChP and 4V ChP (Fig 3G-J).

**Fig 3.**
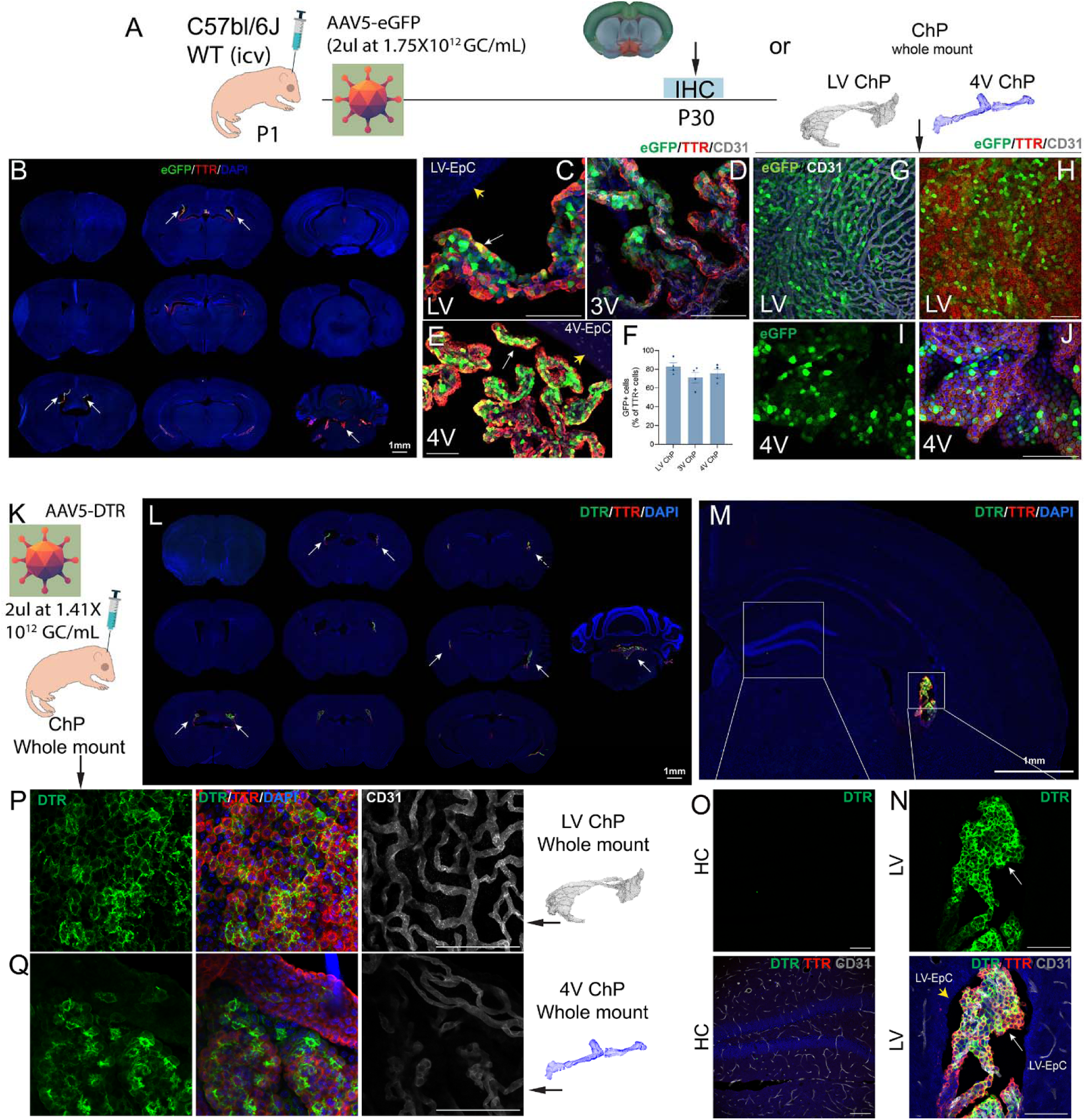
AAV5 CMV-eGFP or DTR show high specificity and efficiency of ChP epithelium transduction in P1 neonates. (A) Experimental design. (B) eGFP expression in the ChP at P30 overlaps with TTR immunostaining. White arrows indicate ChPs in different ventricles. (C-E) Higher-magnification representative images of LV, 3V and 4V ChP showing eGFP expression in TTR+ cells in ChP. Note the absence of eGFP expression in adjacent ependymal cells (yellow arrow). (F) Quantification of % of TTR+ cells that are eGFP+ in ChP in LV, 3V and 4V. Mean±SEM. Each data point is the average of data from one animal. (G-J) Whole-mount LV ChP and 4V ChP show high transduction efficiency and colocalization with TTR immunoreactivity. (K) AAV5 CMV-DTR virus was injected into P1 pups, and mouse brains were collected at P30. (L) Whole brain section scan of P30 brains immunostained for DTR (localized to the plasma membrane in contrast to eGFP) and TTR immunostaining. White arrows indicate ChPs in different ventricles. (M-O) showing higher magnification images of LV ChP with colocalized DTR and TTR immunostaining in ChP (white arrows) but negative in the adjacent hippocampus, cortex or ependymal cells (yellow arrow). (P) Whole-mount ChP in LV or (Q) in 4V showing colocalization of DTR and TTR-immunostained cells that are not colocalized with endothelial cells in ChP (CD31). Scale bar, 100 µm or as indicated.

Based on this result, we produced the AAV5-CMV-DTR virus and injected neonatal P1 pups at a similar titer. ChP whole-mount preparations were harvested at day 30 post-injection (Fig 3K), and consistent with the AAV-eGFP reporter vector, we observed minimal DTR staining outside the ChP (Fig 3L). In contrast, we show highly expressed DTR protein in the ChP epithelial cells in the lateral ventricle (Fig 3M-3N), colocalizing with TTR immunoreactivity, with no or minimal detection of DTR-expressing cells in the adjacent hippocampal, cortical, or ependymal cells (Fig 3M and 3O). Consistently, in the ChP whole mount, DTR immunostaining is highly abundant, localized on the membrane and cytoplasm, and colocalizes with TTR+ CPECs, but is not with endothelial cells (CD31+) either in LV ChP whole mount (Fig 3P) or 4V ChP whole mount (Fig 3Q).

We then evaluated the transduction efficiency and specificity of both AAV5 CMV-eGFP and AAV5 CMV-DTR in young adult mice. We injected the virus at (1.26 × 10^10 GC/mouse) into the left lateral ventricle and examined the expression of eGFP in ChP whole-mount preparations 30 days later. We find similar high efficiency transduction of ChP epithelial cells and high ChP specificity in the adult mouse brain in both ChP whole mounts (Fig 4A-E) and IHC-processed coronal brain sections. Overall, this demonstrates a high transduction efficiency in ChP epithelial cells with minimal or absent transduction in adjacent ependymal cells, hippocampal or cortical tissues (Fig 4F-K). Consistently, in adult mice, AAV5 CMV-DTR virus (1.21 × 10^10 GC/mouse) also yielded highly efficient CPEC transduction with minimal expression in other brain regions, demonstrated both by ChP whole-mount and coronal brain section immunostaining (Fig 4L-U).

**Fig 4.**
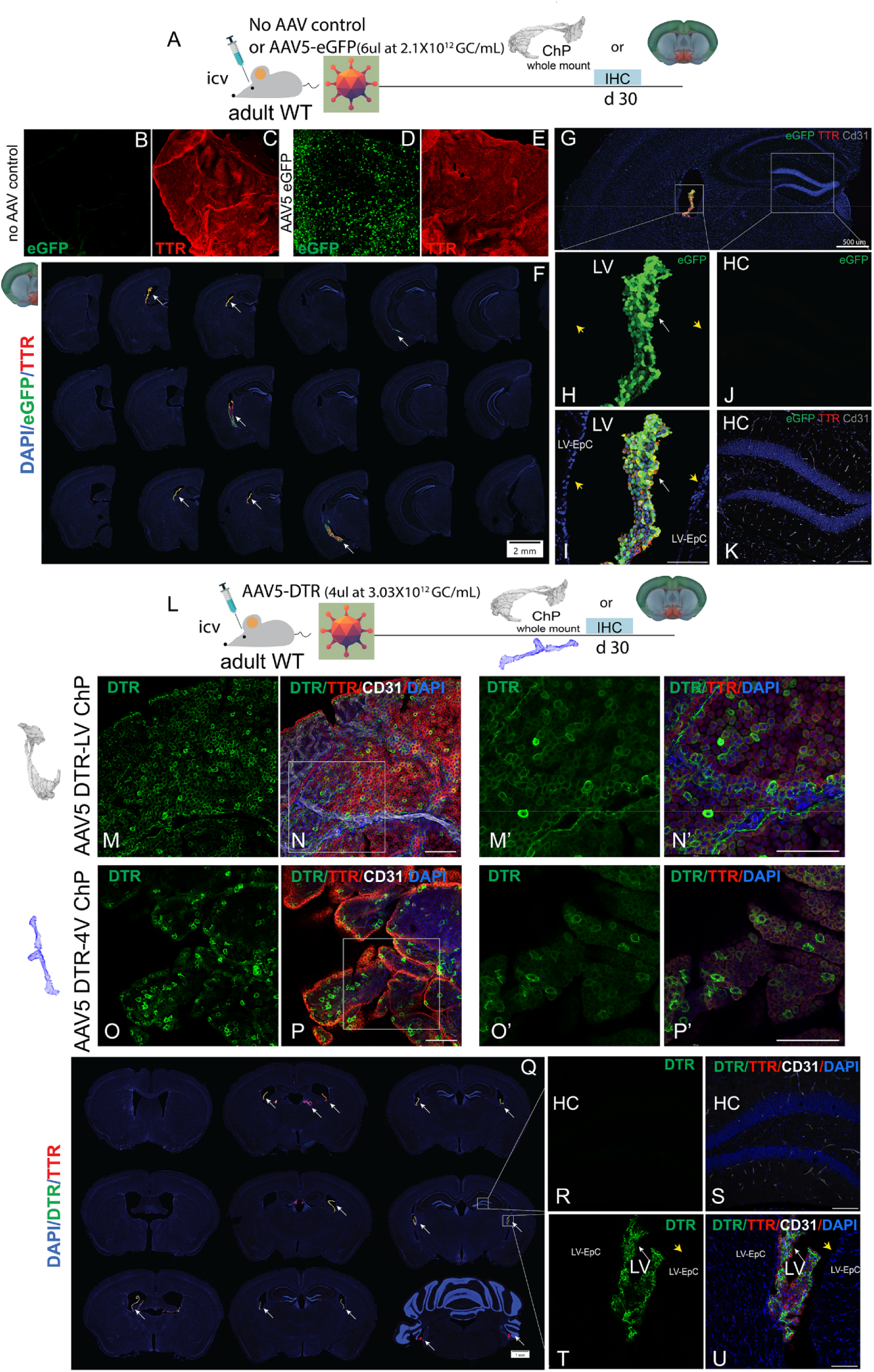
AAV5 CMV-eGFP or DTR show high specificity and efficiency of ChP epithelium transduction in young adult mice. (A) Experimental timeline. (B-C) Whole-mount LV ChP visualized for eGFP expression and TTR immunostaining in mice injected with no AAV control or (D-E) AAV5 CMV-eGFP. (F) eGFP expression in the ChP at d30 overlaps with TTR immunostaining. White arrows indicate ChPs in different ventricles. (G-K) Higher-magnification images of LV ChP with colocalized eGFP and TTR immunostaining in ChP but negative in the adjacent ependymal cells (yellow arrow), hippocampus, and cortex. (L) AAV5 CMV-DTR virus is injected unilaterally into the lateral ventricle of young adult mice and harvested at d30. (M-N’) Whole-mount ChP in LV showing colocalization of DTR and TTR-immunopositive cells that are not colocalized with endothelial cells in ChP (CD31). (O-P’) Colocalization of DTR and TTR immunostaining in 4V ChP whole mount and lack of colocalization of DTR and CD31 immunoreactivity. (Q) DTR expression in the ChP at d30 overlaps with TTR immunostaining. White arrows indicate ChPs in different ventricles. (R-U) Higher-magnification images of LV ChP with colocalized DTR and TTR immunostaining in ChP but negative in the adjacent ependymal cells (yellow arrow), hippocampus, and cortex. Scale bar, 100 µm or as indicated.

With confirmed transduction efficiency and high specificity of the AAV5 serotype in the ChP in all ventricles following a single unilateral injection in neonates (P1) or adult mice, we then evaluated whether delivering DTR in this manner could be a useful tool for ChP ablation to achieve CSF volume loss. We injected neonates at P1 unilaterally with 2 µL of AAV5 CMV-eGFP or CMV-DTR, and administered Dtx subcutaneously daily (20ng/g) for 3 consecutive days on P4-P6 (Fig 5A). AAV-eGFP or DTR-injected pups were subjected to the same Dtx treatment and underwent MRI on P30 to measure CSF volume, and brain tissues were harvested to evaluate ChP tissue loss. We observed substantial loss of TTR+ ChP tissue in AAV-DTR-injected mice compared to AAV-eGFP controls (Fig. 5B-C). Consistently, we observed a profound decrease in CSF volume in the DTR-injected mice. At P30, the final CSF volume was quantified using fluid-sensitive MRI and the 3D ventricular volume reconstructed in Imaris as previously described^31^. Our data show that this experimental paradigm efficiently reduced CSF volume in neonates to a similar extent as in iDTR mice treated at P10 or later (Fig 5D-F). Given our previous study that shows a significant reduction of CSF volume after a single 40 ng/g dose of Dtx in adult iDTR mice with robust DTR expression in the ChP, we speculate that loss of CSF could occur as early as post-single dose at P5 in neonatally treated mice.

Next, we evaluated the effects of neonatal ChP ablation on early postnatal neurogenesis and general health and locomotor function. Wildtype C57Bl/6J neonates were injected at P1 with AAV-eGFP or AAV-DTR and treated with Dtx (20ng/g) or saline (as DTR only control) daily subcutaneously from P4-P6. Brain tissue was then harvested at 30 days of age (Fig. 6 A). We previously reported that loss of ChP and CSF in adult mice starting from 1 month post-Dtx leads to a significantly decreased DCX+ neuroblast pool in the SVZ^31^. Indeed, we replicated this finding in neonatal mice using SVZ whole-mount preparations (Fig 6B-C), suggesting that in early postnatal days, ChP-derived factors or the CSF itself are also critical to maintain the DCX+ NB population in the SVZ (Fig. 6 B, C). Note that AAV-DTR+saline mice (Fig 6C, red points) show similar DCX+ cell density as the AAV-eGFP control (Fig 6C, blue points), supporting that the decreased neuroblast cell pool at the SVZ wall is due to DTR-dependent Dtx-induced ablation of CPECs (Fig 6C, orange points), not the expression of DTR in the CPECs alone. Consistently, coronal section immunostaining shows similar results as the SVZ whole mount (Fig 6D-F). Interestingly, we also observed a different distribution pattern of the DCX+ neuroblasts on the SVZ whole mount, with AAV-DTR+Dtx mice showing isolated clusters of neuroblasts in contrast to the expected net-like pattern reflecting the migratory NB network in control mice (Fig 6B, 6G). Additionally, to evaluate the integrity of ependymal cells in ChP-ablated mice, we stained for an ependymal cell marker (β-catenin). Interestingly, while there is no loss of ependymal cell coverage on the SVZ wall in ChP-ablated mice, the ependymal cell distribution is altered around the neuroblast clusters in ChP-ablated mice, in contrast to the uniform distribution in the control mice (Fig 6G). We next examined postnatal neurogenesis in the SGZ (Fig. 7). It is known that the population of DCX+ NBs sharply decreases with age^26,37^, being highly abundant in the hippocampus of young P30 mice. To ensure that all NBs are quantified despite their high number and close proximity, we used a machine learning approach to quantify SGZ DCX+ cells. Our data show that there is a moderate decrease in the total number of DCX+ cells in the AAV5 DTR ChP-ablated mice (Fig 7B, C, and F), supporting a critical role of ChP-derived factors or CSF in postnatal SGZ neurogenesis as well. To investigate whether the decrease in DCX+ cells is due to the decreased proliferation of neural progenitor cells or increased apoptotic cells in the SGZ, we quantified Ki67+ cell numbers, as well as TUNEL+ apoptotic cells. Surprisingly, both Ki67+ cell number and TUNEL+ cell number were similar between control and ChP-ablated mice (Fig. 7D, F and H), suggesting that the decreased number of DCX+ cells could reflect a difference in neuroblast differentiation. An alternative explanation could be that the peak of apoptosis in ChP-ablated mice could have already passed by P30, as we only analyzed a single time point as a snapshot of ongoing neurogenesis, which is age-dependent and modulated by ChP-secreted or CSF-borne factors which could act not only on NBs, but also on age-dependent NSC proliferative activity^16^. Additionally, comparing AAV-DTR ChP-ablated mice with control mice that received DTR virus but only saline treatment shows similar results (Fig S3), supporting that ChP ablation drives the decrease in hippocampal neurogenesis, and not the expression of DTR in the CPEC. Consistent with the decrease in DCX+ cell numbers, the hippocampal volume in the ChP-ablated mice is also significantly decreased (Fig. 6E and 6G), as well as total brain volume measured by MRI on both P21 and P30 (Fig 6J). We did not observe an apparent effect of sex in our experiment, but future studies with large cohorts of both sexes are needed to confirm this. Neonatal ChP-ablated mice do not show overt deficits in general health and have the same body weight as control mice at P30 (Fig 8A). Open field test (OFT) shows that ChP-ablated mice display a moderate decrease in horizontal activity, movement time, vertical activity, and stereotypic behavior without affecting their average speed in the OFT (Fig 8B-F), suggesting potential mild effects of neonatal ChP ablation on novelty-induced exploratory locomotion without general motor function impairment.

**Fig 5.**
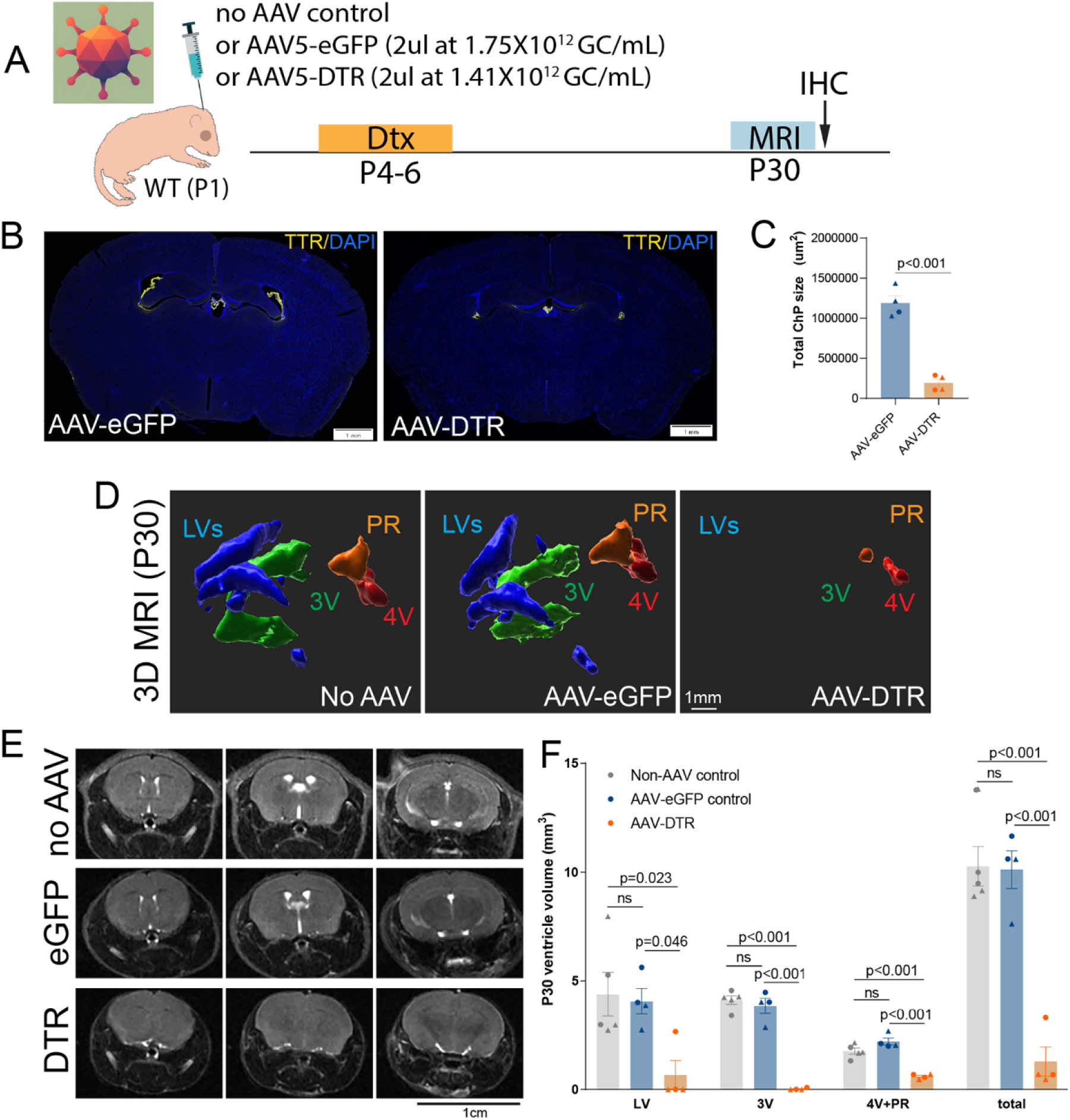
AAV5 CMV-DTR reduces CSF volume in neonatal brains. (A) Experimental timeline and viral dosage in neonatal mice. (B) IHC evaluation of ChP tissue loss using TTR as a marker for CPEC. (C) Quantification of total ChP size (TTR+ area) in experimental groups. (D) Representative fluid-sensitive 3D MRI for all experimental groups and (E) representative T2-weighted MRI scans of all experimental groups showing reduction of all ventricles in AAV5 CMV-DTR injected mice at P30. (F) Quantification of 3D volume of LV, 3V and 4V+PR in experimental groups. Mean±SEM. Each data point is the data from one animal. Circle= female and Triangle=male. *p-values* as indicated, Student’s t-test for panel C and one-way ANOVA (parametric or nonparametric) within each ventricle for panel F.

**Fig 6.**
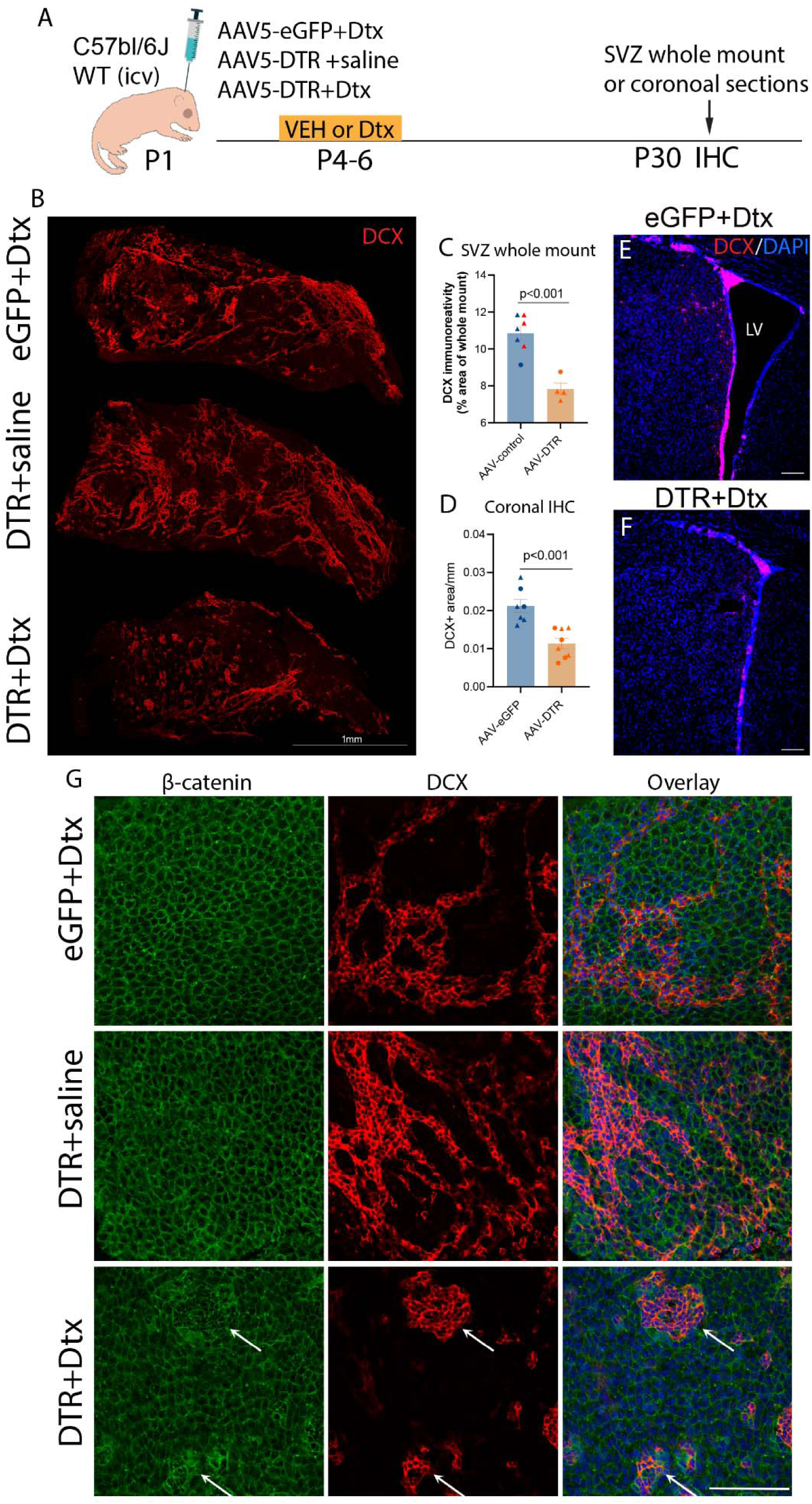
AAV5-DTR-mediated CSF reduction leads to decreased postnatal neurogenesis in the SVZ. (A) Experimental timeline. (B) Representative immunostaining showing DCX (red) in SVZ whole mount in AAV5-eGFP+Dtx, AAV5-DTR+saline, and AAV5-DTR+ Dtx injected mice. (C) Quantification of DCX+ cell density in control (blue dots indicate AAV-eGFP +Dtx control and red dots indicate AAV-DTR+saline control) or AAV-DTR+Dtx (orange bar and dots) mice. (D) Quantification of DCX+ cells/mm length in coronal sections and (E-F) Representative immunostaining showing DCX (red) and DAPI (blue) at SVZ in AAV5-eGFP and AAV5-DTR+Dtx-injected mice. (G) High-magnification images of the ependymal layer (β-catenin, green) and underlying DCX+ (red) neuroblast cell network in control or ChP-ablated SVZ whole mounts. White arrow depicts disorganized ependymal cells on top of the DCX+ clusters in ChP-ablated brains. Mean±SEM. Each data point is the average data from one animal. Circle=female and Triangle=male. *p-* values as indicated; Student’s t-test for panels C and D.

**Fig 7.**
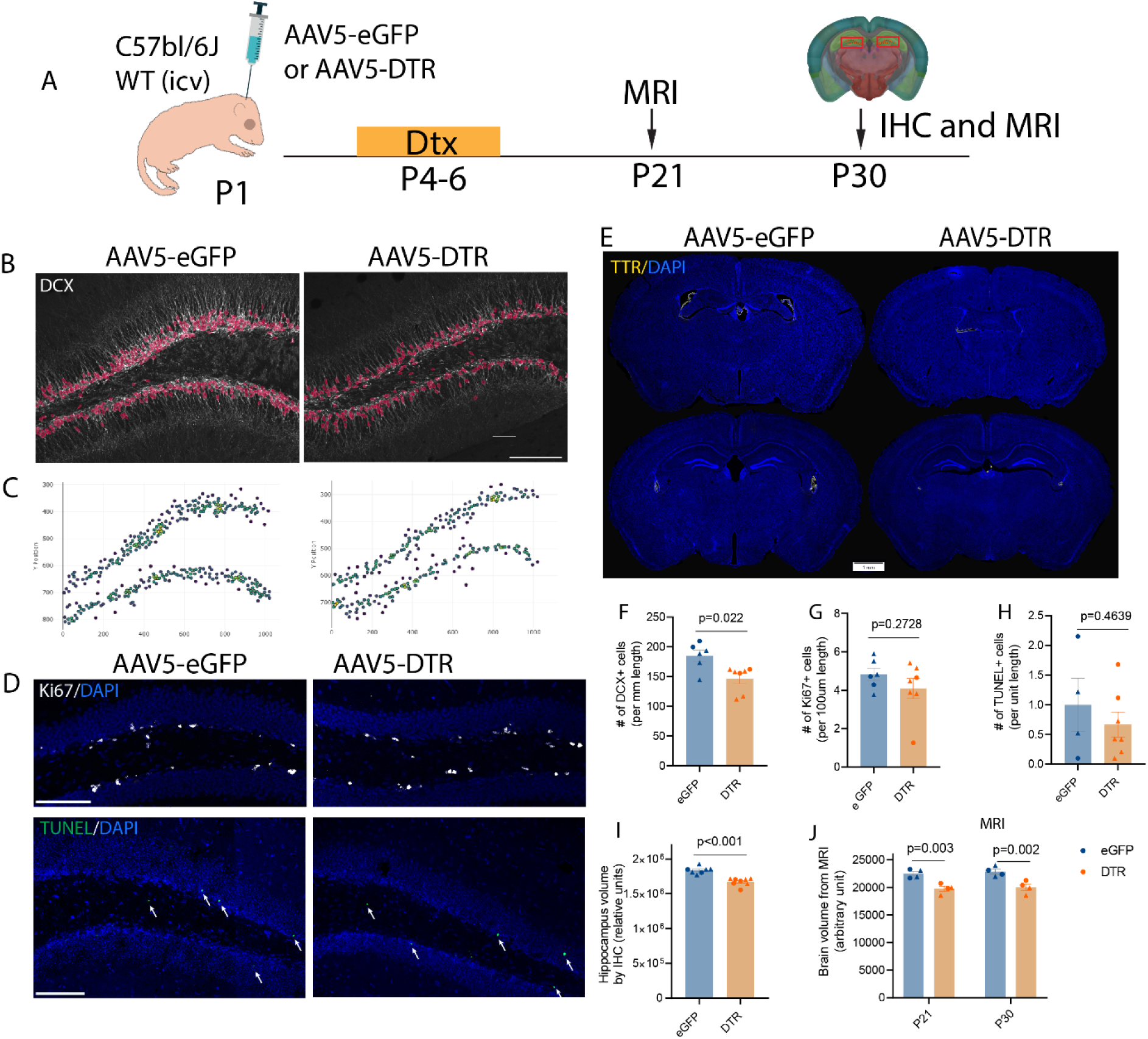
AAV5-DTR-mediated CSF reduction leads to decreased postnatal neurogenesis in the SGZ. (A) Experimental timeline. (B) DCX immunostaining superimposed with AI-model-recognized DCX+ cells and (C) AI-model-recognized DCX+ cells and their coordinates in SGZ. (D) Representative immunostaining showing Ki67 (gray) and DAPI (blue) or TUNEL+ cells (green) at SGZ in AAV5-eGFP and AAV5-DTR injected mice at P30. (E) Representative DAPI staining of the whole brain section of AAV-eGFP or AAV-DTR-injected mouse brains containing the hippocampus. (F-H) Quantification of DCX+, Ki67+, and TUNEL+ cells per unit length of SGZ. (I) Quantification of hippocampal volume (from 6 consecutive hippocampal sections) in AAV-eGFP or AAV-DTR-injected mice. (J) Quantification of whole brain volume using MRI T2-weighted brain sections (12 sections) at P21 and P30. Mean±SEM. Each data point is the average data from an individual animal. Circle=female and Triangle=male. *p-values* as indicated, Mann-Whitney U test for panels F and I, Student’s t-test for panels G and H, Two-way ANOVA for panel J.

**Fig 8.**
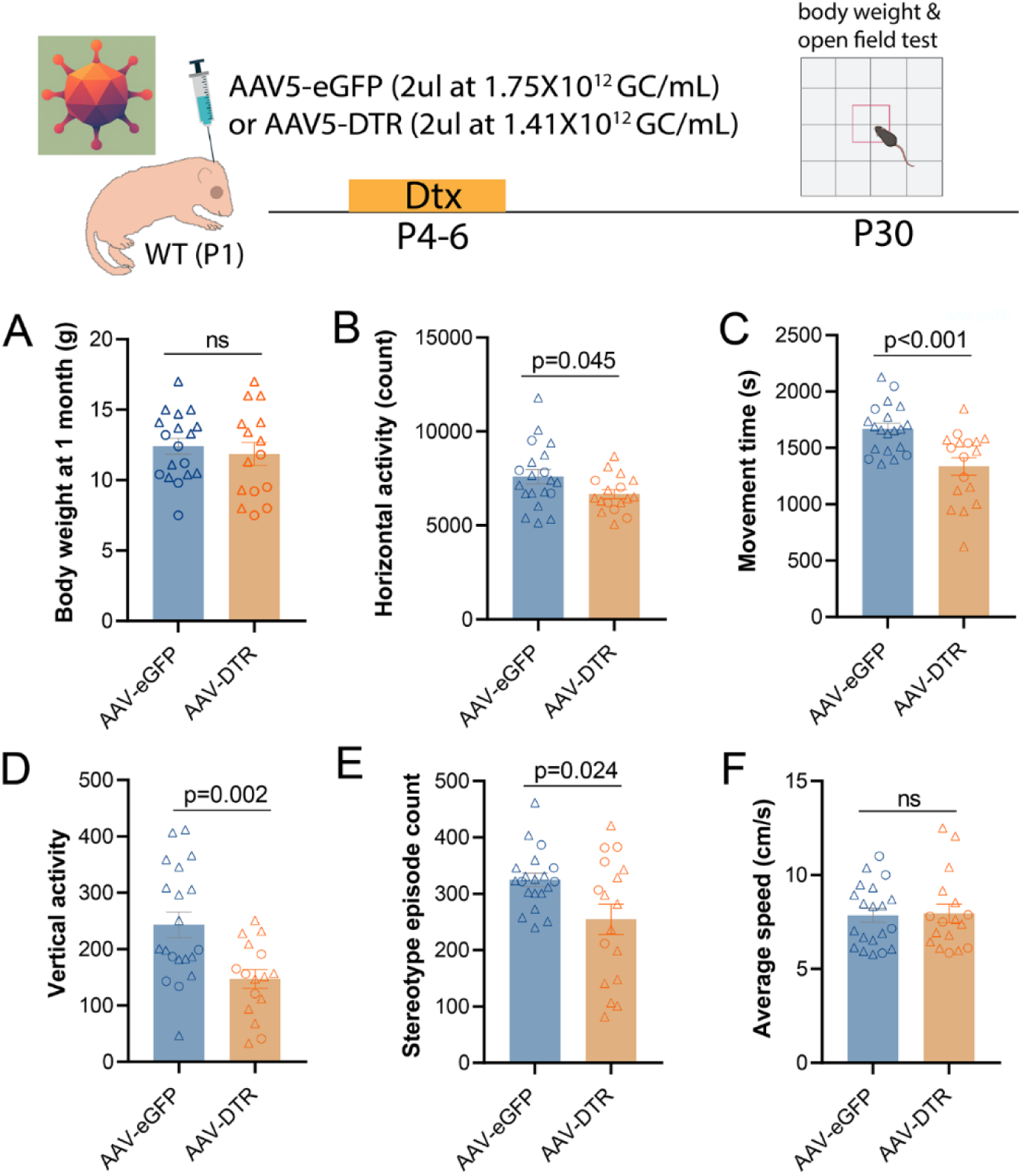
Neonatal ChP ablation decreases novelty-induced exploration in OFT at P30. (A) Experimental timeline. (B) Body weight at P30. (C) Horizontal activity (count), (D) Movement time (s), (E) Vertical activity, (F) Stereotype episode count, and (G) Average speed (cm/s) of the first hour in the open field test chambers. Mean±SEM. Each data point is the data from an individual animal. Circle=female and Triangle=male.. *p-values* as indicated, Student’s t-test for panel A, Welch’s t-test for panels B and E, Student’s t-test for panels C, D and F.

Lastly, we investigated the potential use of AAV5-DTR in hydrocephalus treatment in an adult kaolin hydrocephalus model. First, we examined the efficacy of AAV5-DTR in ablating the ChP of the adult mouse brain. Adult mice (3 to 6 month-old) were injected unilaterally with 1.2 × 10^10^ viral particles of either CMV-eGFP or CMV-DTR and received Dtx injections (20ng/g) daily i.p. on post-surgery days 12-14 and subjected to T2 MRI on post-surgery day 20 (Fig 9A). Brain tissues were harvested to evaluate ChP tissue loss after MRI scan. We observed substantial loss of TTR+ ChP tissue in AAV-DTR injected mice compared to AAV-eGFP controls (Fig. 9B-C). MRI shows substantial loss of ventricular volume (Fig 9D-F), to a similar extent as seen in the adult iDTR mouse model. Therefore, our data show that the AAV5 CMV-DTR viral vector is effective in inducing ChP ablation and CSF reduction in adult mice as well. Given that AAV-mediated ChP ablation and the resulting reduction in CSF volume are not associated with mortality and do not impair motor or cognitive function in mice^31^, we evaluated the therapeutic efficacy of AAV5-CMV-DTR in the treatment of hydrocephalus. Kaolin-induced hydrocephalus is a well-established experimental model in which intracisternal injection of kaolin elicits an inflammatory response that obstructs CSF flow and impairs CSF absorption, resulting in progressive ventriculomegaly and elevated intracranial pressure^59–62^. We first induced DTR expression in the ChP epithelial cells of C57BL/6J mice by intraventricular injection of AAV5-CMV-DTR. Control mice received AAV5-CMV-eGFP. To this end, we utilized the kaolin model as a well-established hydrocephalus paradigm that induces ventricular expansion and parenchymal edema characteristic of hydrocephalus pathology seen in patients. At 10 days post AAV injection, hydrocephalus was induced by intracisternal injection of kaolin, and 6-10 hours after Dtx were administered once daily for 3 consecutive days (20 ng/g/day X3) to ablate the ChP (Fig. 10 A). T2 and 3D fluid-sensitive MRI performed on day 4 and 7 post kaolin injection showed the presence of ventriculomegaly and severe parenchymal edema that expanded from day 4 to day 7 in control mice (Fig. 10B-I). Ablation of the ChP markedly reduced kaolin-induced parenchymal edema (Fig. 10B-I) and white matter edema (measured in the corpus callosum, Fig 10H-I) without reducing ventriculomegaly (Fig. 10F) at both time points, with no expansion of parenchymal edema or white matter edema in ChP-ablated hydrocephalus mice. We conclude that ChP contributes to transependymal CSF flow and the development of parenchymal edema during the early stages of hydrocephalus, and that ChP ablation may represent a promising therapeutic strategy for hydrocephalus.

**Fig 9.**
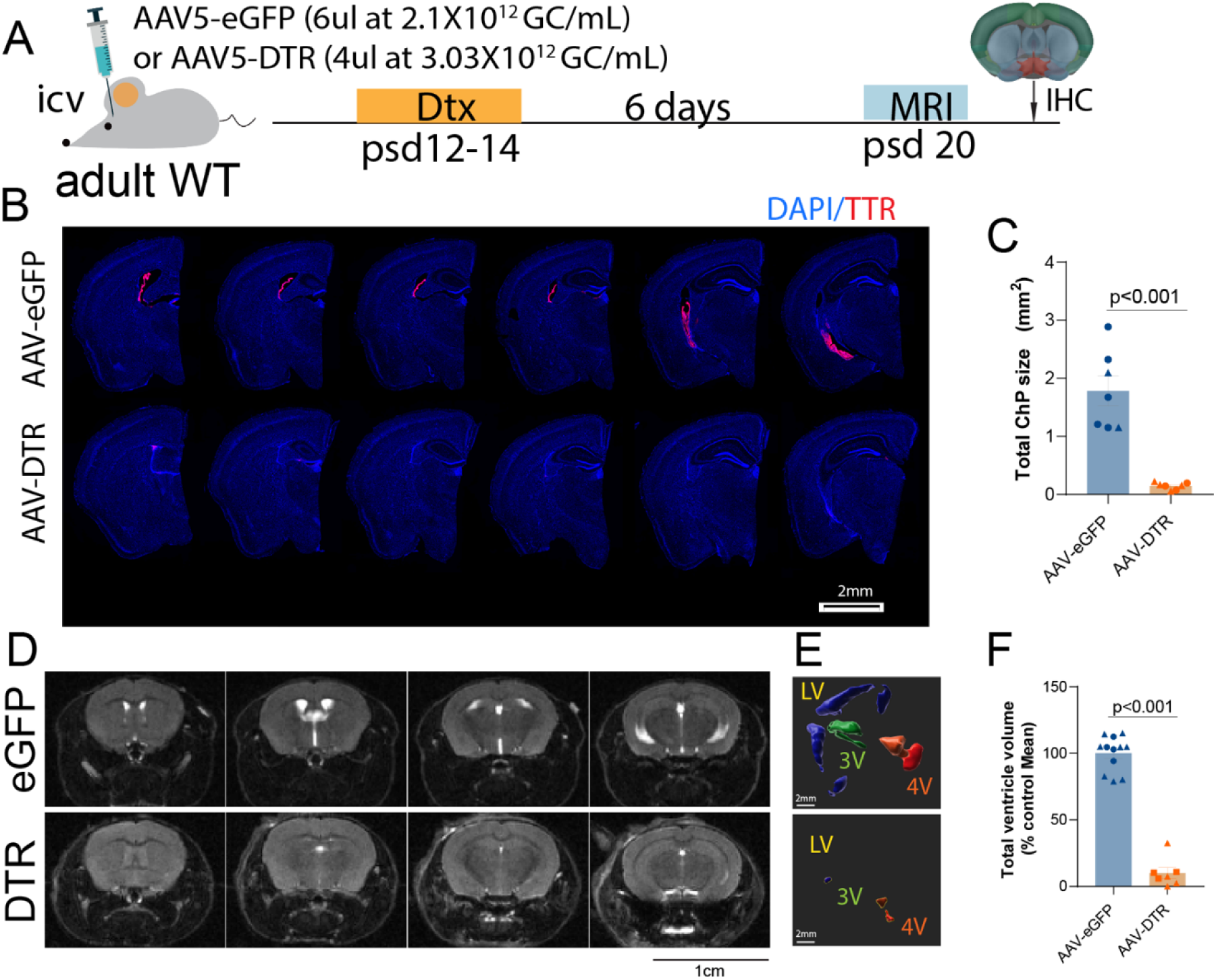
AAV5 CMV-DTR reduces CSF volume in adult brains. (A) Experimental timeline and viral dosage in adult mice. (B) IHC evaluation of ChP tissue loss using TTR as a marker for CPEC. (C) Quantification of total ChP size (TTR+ area) in experimental groups. (D) Representative fluid-sensitive T2-weighted MRI scans for all experimental groups and (E) Representative 3D MRI Imaris reconstruction of all experimental groups showing reduction of all ventricles in AAV5 CMV-DTR injected mice. (F) Total ventricle volume was quantified from a series of T2-weighted MRI scans (7 sections quantified per mouse). Mean±SEM. Each data point is the data from one animal. Circle=female and Triangle=male. *p-* values as indicated, Welch’s t-test for panel C, and Mann-Whitney U test for panel F.

**Figure 10.**
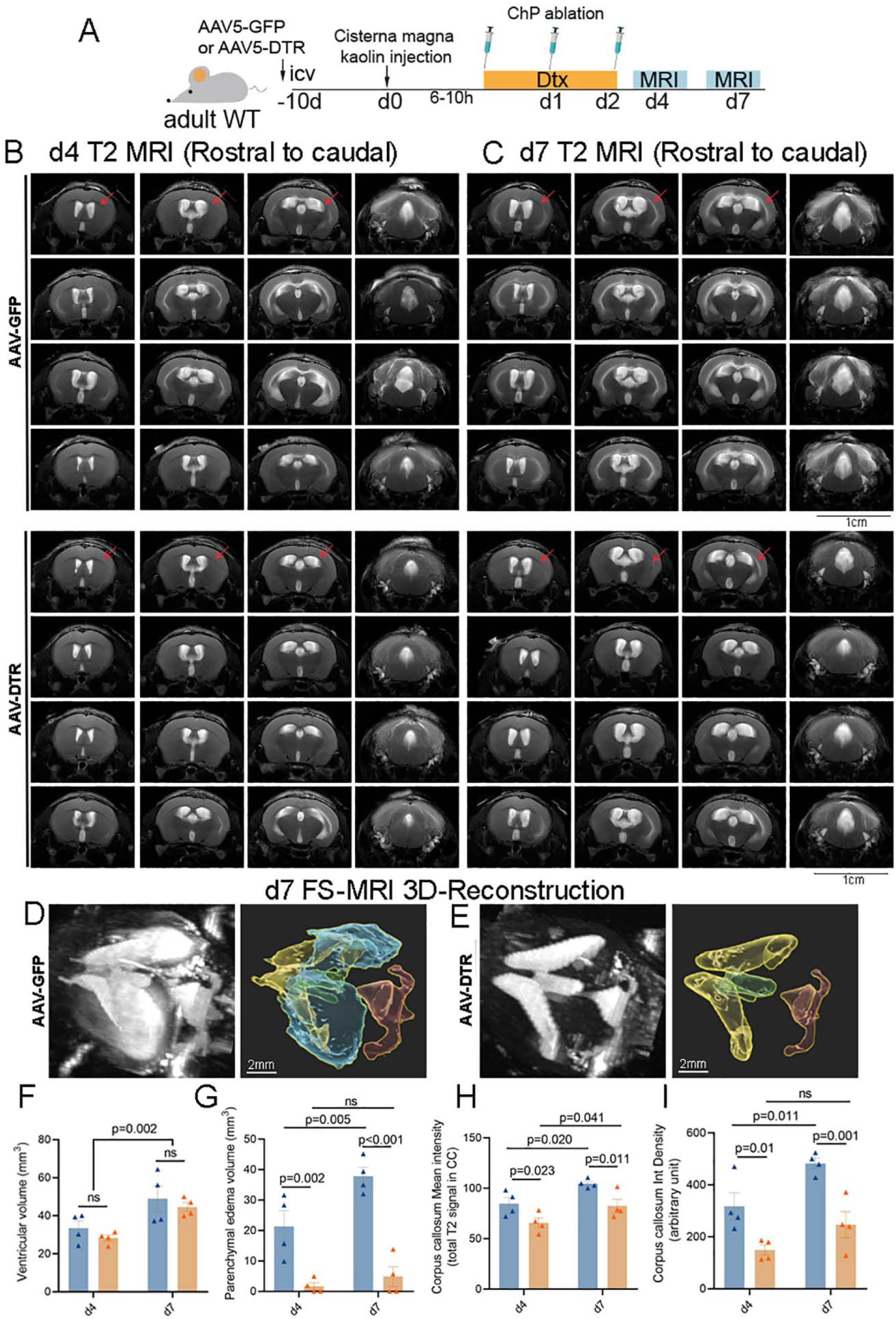
Choroid plexus ablation via AAV-DTR ameliorates hydrocephalus pathology without causing ventricular collapse in adult mice. (A) Experimental timeline. (B, C) T2 MRI slices from kaolin-treated AAV-eGFP (B) and AAV-DTR (C) animals at d4 or d7 post-kaolin injection. Red arrows indicate corpus callosum regions where hyperintensity is seen after kaolin administration. (D-E) Representative 3D volume rendering of MRI T2 hyper-intense regions (left) and segmented 3D objects (right; LV – yellow; 3V – green; 4V – red; parenchymal edema – blue). (F) Quantification of total ventricle volume and (G) parenchymal edema volume from 3D MRI reconstructions. (H) Quantification of mean signal intensity and (I) optical density in the corpus callosum of T2 MRI slices (B). Mean±SEM. Each data point is the average data from one animal. *p-values* as indicated. Two-way ANOVA for panels F-I.

## Discussion

### ChP/CSF loss in neonates decreases SVZ and SGZ neurogenesis, accompanied by moderately reduced exploratory locomotion at P30

In our prior work, we demonstrated that ChP ablation and CSF loss in young adult ROSA26-iDTR mice markedly reduced the pool of SVZ DCX+ neuroblasts by promoting their premature migration into the olfactory bulb, while leaving overall proliferative activity in the SVZ intact^31^. Here, we extend these findings to the hippocampal neurogenic niche and show that ChP/CSF loss in adolescent iDTR mice leads to a moderate reduction in SGZ DCX+ neuroblasts without changing the number of Ki67+ proliferating cells. Similarly, in neonatal wild-type mice subjected to AAV5-mediated ChP ablation and CSF volume loss following Dtx treatment at P3-5, we observe a decrease in both SVZ and SGZ DCX+ cells and a reduction in hippocampal and overall brain volume at P30, again without a difference in Ki67+ TUNEL-positive apoptotic cell numbers at this time point.

These results suggest that ChP/CSF is not required to sustain the overall proliferative pool in the SGZ, but instead contributes to the differentiation, maturation, and/or survival of newly generated neuroblasts, albeit a peak of apoptosis induced by ChP ablation that occurs prior to P30 cannot be excluded. The modest magnitude of the effect on the SGZ NB pool, compared with the pronounced SVZ phenotype we previously reported, likely is due to the closer contact of the SVZ with the CSF in LVs, as well as the dependence of SVZ NB migration on the CSF flow generated by ependymal cilia on the SVZ wall^11^, whereas the SGZ is localized within the parenchyma of the dentate gyrus, and thus more distally from the lumina of the LV and 3V. Moreover, SGZ NBs migrate only within the GCL, suggesting different chemotactic mechanisms regulating this migration. Interestingly, we also observed disorganized ependymal cell morphology in the SVZ wall of ChP-ablated animals in whole-mount preparations. Such disorganized ependymal cells were associated with the isolated neuroblast clusters in ChP-ablated animals. Although we did not observe loss of ependymal coverage in the lateral ventricle walls of ChP-ablated animals, this colocalization of disorganized ependymal cells with the neuroblast clusters supports close crosstalk and interaction of ependymal cells with underlying neuroblasts in the SVZ. Whether aberrant neuroblast or ependymal cell disorganization drives the other component warrants further investigation in future studies. Nonetheless, our data indicate that ChP/CSF are needed to maintain the NB pool both in the SGZ and SVZ during the postnatal period. As SGZ neurogenesis has not been previously considered to be regulated by ChP/CSF, our study opens new avenues to modulate SGZ neurogenesis through targeting ChP-secreted or other CSF-borne factors to treat diseases characterized by reduced hippocampal neurogenesis, such as aging^2^, Alzheimer’s disease^26^, depression^24^, Parkinson’s disease^63,64^, and type 2 diabetes^65,66^. While our previous study showed that ChP-ablation and CSF loss in adult mice do not affect general motor and cognitive function, this study shows reduced exploratory locomotion in the OFT in ChP-ablated mice, suggesting that ChP ablation during the neonatal period might have more severe side effects compared to adulthood.

### Age dependence of ROSA26-iDTR-mediated ChP ablation and development of AAV5-DTR as a complementary tool

This study reports the age dependence of ROSA26-iDTR-mediated ChP ablation. While adolescent and adult iDTR mice exhibit robust (>90%) loss of CSF volume after Dtx treatment, recapitulating our previous results, the same Dtx regimen applied at P3-5 produces minimal reduction in ventricular CSF volume. Even doubling the Dtx dose (40ng/g daily) at P3-5 yields only modest reductions in CSF volume. Additionally, we observed considerable Dtx toxicity resulting in high mortality at doses higher than 20 ng/g/dayX3, making even modest CSF volume loss induction impractical in iDTR pups and necessitating the use of Dtx-treated controls to avoid confounding effects. In contrast, Dtx administration beginning at P10-12 achieves an efficiency of ChP/CSF ablation comparable to that seen in older animals.

These data suggest that either the “leaky” DTR expression in ChP epithelial cells or the susceptibility of these cells to Dtx-mediated ablation is developmentally regulated and lowest at early postnatal stages. Given that AAV-DTR-transduced neonates ablate successfully when Dtx is delivered during P4-6, it suggests that ChP epithelial cells are susceptible to Dtx-induced apoptosis when DTR is expressed in abundance. We speculate that the “leaky” DTR expression is not abundant until approximately P10 in the ROSA26-iDTR mice. As a result, this mouse model is a robust tool for manipulating ChP/CSF in mice older than P10 but is poorly suited for modeling neonatal conditions that require early ChP manipulation, such as neonatal hydrocephalus or perinatal brain injury.

To overcome this limitation, we developed an AAV5-based strategy for ChP-specific expression of DTR. Building on our prior work showing that AAV5 efficiently transduces ChP epithelial cells^49,50,58^, we demonstrate that AAV5-CMV-eGFP, delivered via unilateral intraventricular injection at P1, transduces ∼70% of transthyretin-positive ChP epithelial cells in lateral, third, and fourth ventricles with minimal off-target expression in other brain regions. AAV5-CMV-DTR exhibits a similarly high degree of efficiency and specificity in both neonatal and adult mice, as shown by robust DTR expression in ChP epithelial cells, and minimal labeling of endothelial or ependymal cells.

When combined with systemic Dtx administration, AAV5-CMV-DTR effectively reduces CSF volume in both neonatal and adult mice to an extent comparable to that seen in ROSA26-iDTR animals treated after P10. In neonates, unilateral AAV5-DTR injection at P1 followed by Dtx treatment at P4-P6 produces significant loss of ChP tissue and ventricular shrinkage by P14 and P30, whereas AAV5-DTR alone does not affect CSF volume, confirming that CSF loss is driven by DTR-dependent ChP ablation. In adults (3–6 months), AAV5-DTR plus Dtx likewise leads to loss of TTR+ ChP tissue and substantial ventricular volume loss, paralleling the phenotype in adult iDTR mice. These results validate AAV5-CMV-DTR as a flexible tool for ChP ablation and CSF volume reduction across a broad age range, including early postnatal stages where iDTR is ineffective. Additionally, since the ROSA26iDTR mouse line carries the loxP-flanked STOP cassette, it makes it difficult to combine this model with other cell-type-specific cre-loxP gene induction or deletion models. The AAV5-DTR vector can circumvent this challenge. This valuable tool could have broad potential usage in investigating the efficacy of ChP ablation in the treatment of hydrocephalus, especially neonatal hydrocephalus, since the iDTR mice do not achieve robust ablation until P10.

### Converging roles of ChP/CSF in SVZ and SGZ neurogenic niches

Our previous work in ROSA26-iDTR mice established that ChP/CSF is dispensable for overall SVZ proliferation but required to retain a pool of newly born SVZ DCX neuroblasts and to support their recruitment to ischemic lesions^31^. The current study extends these insights to the hippocampal SGZ, revealing that ChP/CSF loss modestly reduces the number of SGZ DCX cells and decreases hippocampal volume during a period of robust postnatal neurogenesis.

Taken together, these findings suggest that ChP/CSF fulfills two related but distinct roles in the postnatal neurogenic niches: In the SVZ, ChP/CSF maintains a reservoir of newly born neuroblasts by restraining premature migration to the olfactory bulb and by supporting injury-induced neuroblast mobilization into damaged regions. In the SGZ, ChP/CSF supports postnatal hippocampal neurogenesis and dentate gyrus growth, likely by modulating differentiation or maturation of neuroblasts and/or adult-born granule neurons rather than by directly influencing NSC proliferation.

The mechanisms underlying these effects may involve both biochemical and biophysical factors. ChP-ablation in both the ROSA26-iDTR mouse model and the AAV-DTR platform leads to reduction of CSF volume and collapsing of the ventricular system, making it difficult to distinguish the effects of volume reduction in CSF or altered composition of CSF. ChP-derived secreted molecules (e.g., growth factors, cytokines, guidance cues) can act directly on NSCs or neuroblasts^12,14^, whereas CSF production or flow may provide structural mechanical forces that may be needed for proper niche architecture and guidance cues. In our previous work, ChP ablation disrupted ependymal cilia bundles^31^, the flow generated by which controls SVZ neuroblast migration^11^, and it remains possible that structural or extracellular matrix component alterations contribute to the hippocampal phenotype. Future studies will be needed to distinguish the relative contributions of specific ChP-derived factors, CSF flow dynamics, and mechanical constraints to SVZ and SGZ neurogenesis.

### AAV-mediated ChP ablation prevents parenchymal edema in hydrocephalus

Our findings demonstrate that ablation of the ChP during the early phase of hydrocephalus, 6-10 hours after disease induction, prevents the development of parenchymal edema. During the initial hours following fourth ventricular outflow obstruction due to the introduction of kaolin into the cisterna magna, continued CSF secretion by the ChP likely increases intraventricular pressure, promotes transependymal CSF movement into the surrounding white matter, and initiates edema formation^44^. Interrupting this process before edema becomes established appears sufficient to preserve parenchymal water homeostasis. Hydrocephalus is a dynamic process in which ventricular enlargement and transependymal CSF flow precede a cascade of secondary pathological events, including white matter injury, neuroinflammation, axonal damage, and disruption of the blood-brain and ependymal barriers. Once these downstream mechanisms are initiated, simply reducing CSF production may no longer be sufficient to reverse tissue injury. Our data therefore identify an early therapeutic window during which targeting the ChP can modify disease progression before irreversible pathological changes would require permanent CSF diversion surgery^43^.

Our data on postnatal SVZ and SGZ neurogenesis showed that early-life ChP dysfunction or CSF depletion may moderately impair the neuroblast pool in the postnatal neurogenic niches, potentially influencing cognitive or affective trajectories later in life. However, given the devastating effects of hydrocephalus on CNS development and function^39^, if ChP ablation partially corrects that, the benefits might still outweigh possible side effects of ChP ablation in neonatal hydrocephalus, while avoiding invasive interventions such as surgical ChP cauterization or endoscopic third ventriculostomy^38,46^. Our novel AAV5 CMV-DTR viral vector demonstrates that AAV-mediated ChP ablation is achievable in neonatal mouse brains as early as P4-6, and the reduction of CSF is stable at least up to P30. Combined with the rescue of hydrocephalus pathology that we observed in the adult kaolin model, these data provide the first evidence and proof of principle that AAV-mediated gene therapy to ablate ChP could be a promising treatment strategy for disorders of CSF homeostasis. Specifically, ChP ablation reduced parenchymal edema and corpus callosum hyperintensity, an indicator of white matter damage, both key features of hydrocephalus pathology^42–45^. Whether AAV5 or other serotypes of AAV could have similar high tropism and efficiency in the human ChP epithelial cells is an aspect that remains to be investigated. Conversely, these results also highlight the need to consider age, timing, and residual ChP/CSF function when designing ChP-targeted therapies in neonates and children.

## Limitations and Future Directions

Several limitations should be noted. First, our analyses of SGZ proliferation and apoptosis are based on a single time point (P30 for neonatal studies, or 3 months post-ablation for adult mice) and may miss transient changes in proliferation or cell death that could occur earlier. BrdU/EdU birthdating and apoptosis assays over multiple time points will be needed to determine how ChP/CSF loss alters the kinetics of NSC proliferation, neuronal differentiation and survival over time. Second, we did not directly assess cognitive function in the current study; thus, the functional consequences of the observed reductions in SGZ neurogenesis and hippocampal volume remain to be established. Third, although AAV5-CMV-DTR shows high specificity for ChP epithelial cells, low levels of off-target transduction in other regions cannot be entirely excluded and should be carefully monitored, particularly when combining this approach with additional genetic manipulations.

We did not examine CSF composition (e.g., ions, proteins, RNAs) or the blood-CSF barrier properties after ablation. Integrating our ablation tools with proteomic and transcriptomic profiling of residual CSF and ChP and brain tissues will be essential to identify candidate factors linking ChP/CSF to SVZ and SGZ neurogenesis. Nevertheless, the present study, together with our earlier work, underscores the role of ChP/CSF in shaping both SVZ and SGZ neurogenic niches and expands the toolkit for ChP ablation, namely ROSA26-iDTR for late juvenile/adult stages and AAV5-DTR for neonatal and adult stages to study ChP/CSF functions in development, injury, and disease. Lastly, future studies comparing early and delayed ChP ablation in other hydrocephalus models would be needed to determine the therapeutic window and optimal timing of ChP-targeted interventions in hydrocephalus. Additionally, histopathological evaluation of neuronal and white matter damage as well as neuroinflammatory response in non-ablated or ChP-ablated hydrocephalus animals will be valuable in providing mechanistic insights into the beneficial neuroimaging findings we observed in the kaolin model and may identify novel therapeutic targets for reducing hydrocephalus-associated pathology and brain tissue damage.

## Authorship contribution statement

AT and YL conceptualized and designed the overall study and performed the majority of the experiments. The hydrocephalus experiment was designed and analyzed by FM, OVC, CS, YL, and AT. FM and OVC carried out the hydrocephalus experiment and acquired the data. AT carried out neonatal AAV viral injection and Dtx treatment. AT, SH, EK and JP carried out IHC staining, imaging, and quantification. AT, SH, JP, OVC, CS and YL drafted and edited the manuscript. FW carried out part of the MRI quantification and OL carried out part of the imaging. LM maintained the mouse colony.

## Consent for publication

Not applicable.

## Ethics approval and consent to participate

All animal procedures were performed in accordance with the Guide for the Care and Use of Laboratory Animals (8th Edition, National Research Council) and approved by the Institutional Animal Care and Use Committee (IACUC) at University of Cincinnati (Protocol Number 24-02-27-01) and University of California, Davis (Protocol Number 23673).

## Acknowledgements

We thank Chet Closson and the University of Cincinnati live imaging core (supported by NIH S10OD030402) for technical support. We thank Diana Lindquist, Elizabeth Fugate, Sharon Wang, and Lisa Lemen at the University of Cincinnati, and Sydney Leigh Huddleston and Brad Hobson at UC Davis for assistance with MR imaging.

## Data availability

### Materials availability

The vectors used in the study can be purchased from VectorBuilder.

### Data and code availability

Microscopy data reported in this paper will be shared by the lead contact upon request. No original code was generated in this study. Any additional information required to reanalyze the data reported in this paper is available from the lead contact upon request. All raw data are provided in an Excel file.

