## Supplemental Figures 1-3 for "Ablation of the choroid plexus attenuates hydrocephalus induced parenchymal edema but moderately reduces postnatal hippocampal neurogenesis"

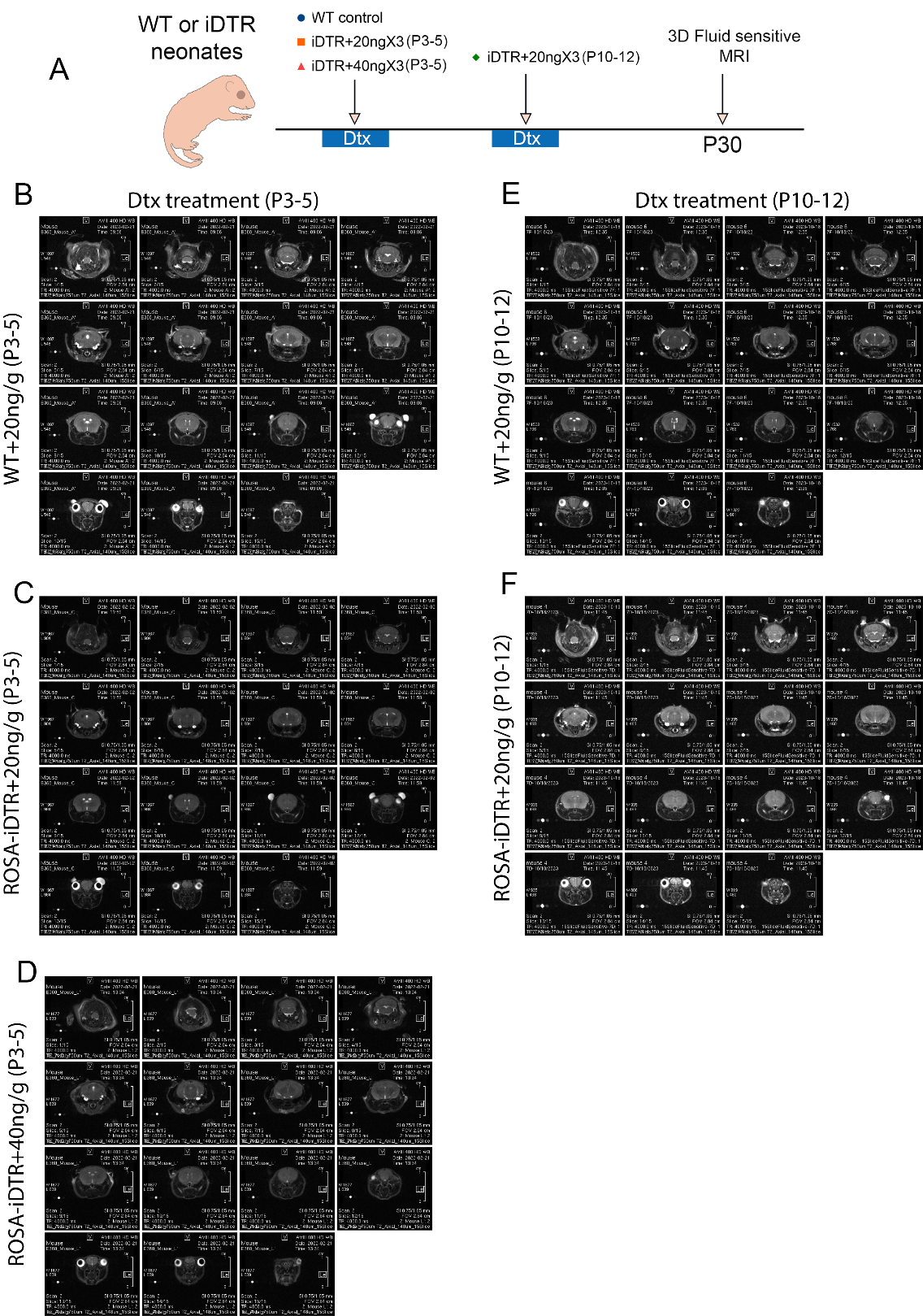

**Supplemental Fig 1.** (A) Experimental timeline. (B-D) Representative T2-Weighted MRI for WT or iDTR neonates treated with 20ng/g daily or 40ng/g daily Dtx at P3-5. (E-F) WT or iDTR neonates treated with 20ng/g daily Dtx at P10-12.

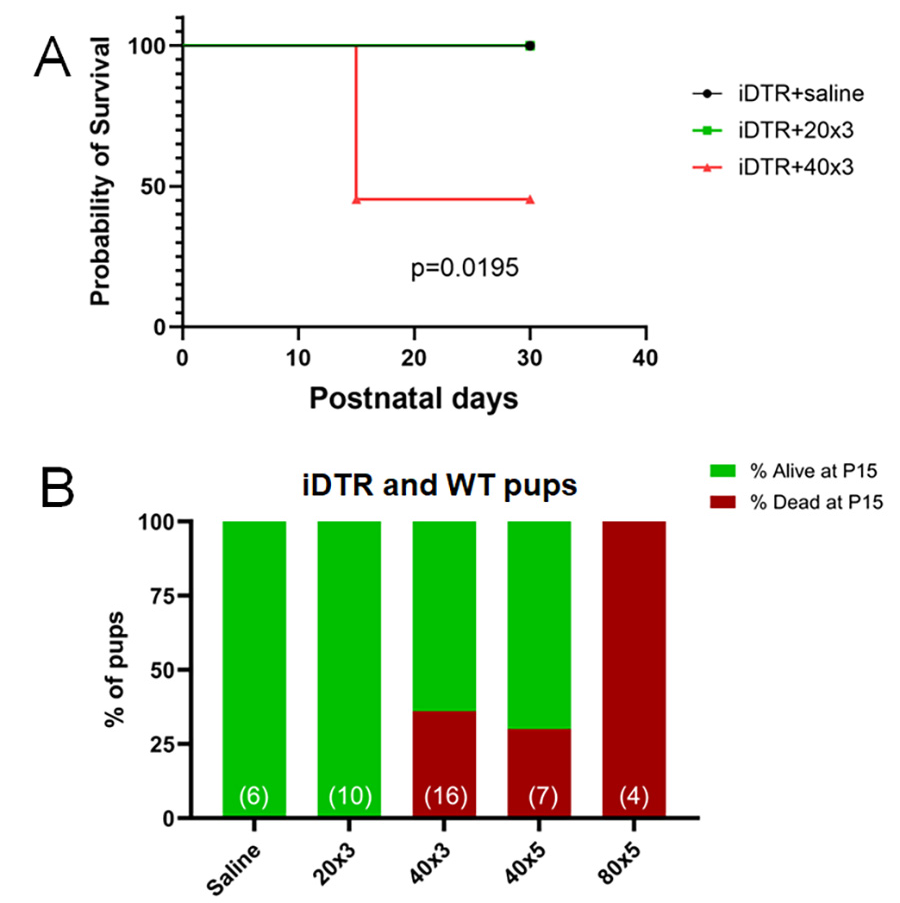

**Supplemental Fig 2.** (A) Survival curve of iDTR neonates that received different dosages of Dtx at P3-5. Log-rank Mantel-Cox test. (B) Percentage of neonates (iDTR and WT) that survived or were dead at P15 following different dosages of P3-5 Dtx treatment. Total number of animals per dosage regimen indicated in parentheses.

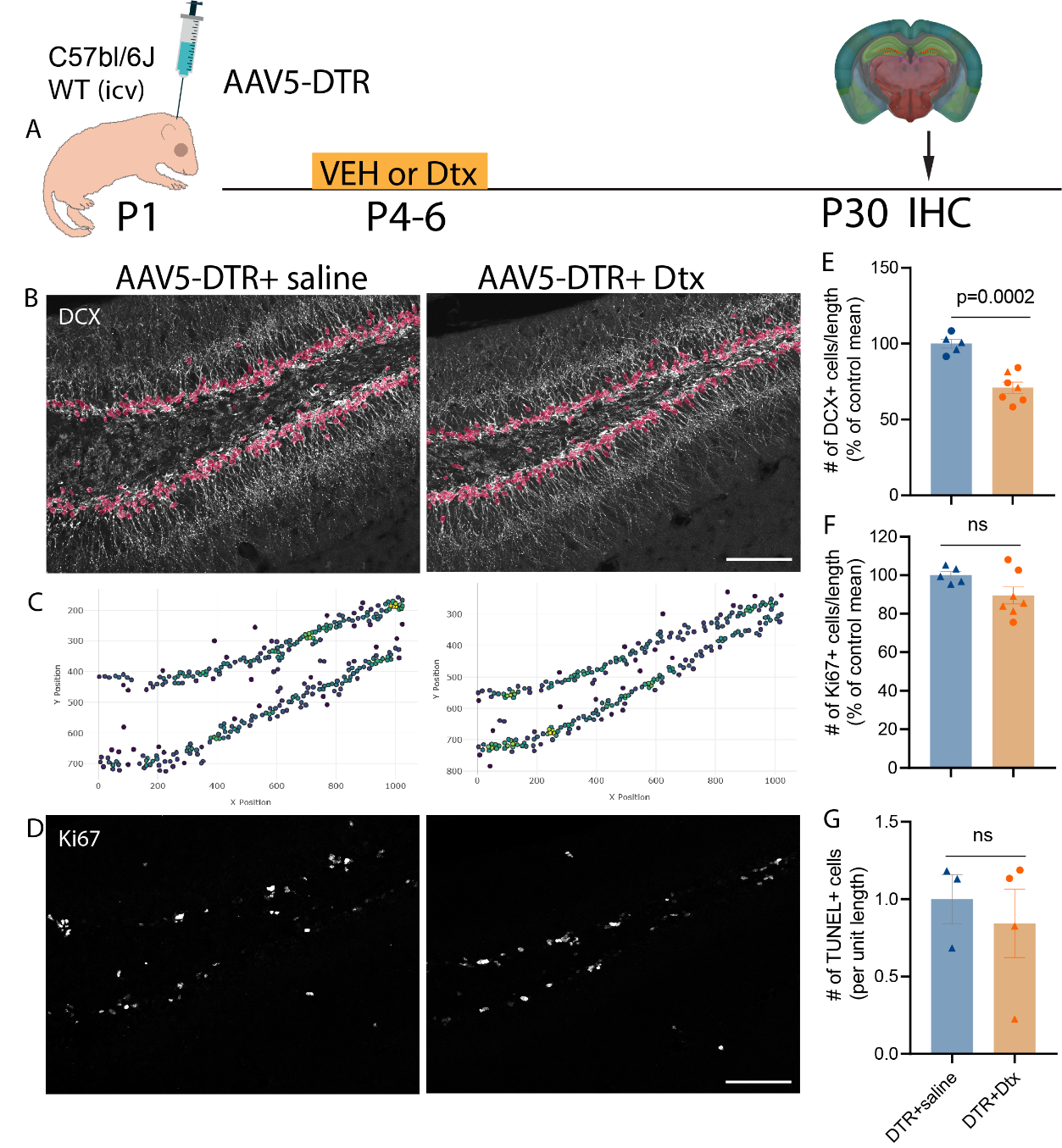

**Supplemental Fig 3.** AAV5-DTR-mediated CSF reduction leads to decreased postnatal neurogenesis in the SGZ when comparing AAV5-DTR injected mice treated with saline or Dtx. (A) Experimental timeline. (B) DCX immunostaining superimposed with AI-model-recognized DCX+ cells and (C ) AI-model-recognized DCX+ cells and their coordinates in SGZ. (D) Representative immunostaining showing Ki67 (gray) at SGZ in AAV5-DTR saline- or Dtx-injected mice at P30. (E-G) Quantification of DCX+, Ki67+ and TUNEL+ cells per unit length of SGZ. Mean±SEM. Each data point is the average data from an individual animal. Circle=female and Triangle=male. *p-*values as indicated, Student’s t-test for panels E, F and G.
